# Theca cell mechanics and tissue pressure regulate mammalian ovarian folliculogenesis

**DOI:** 10.1101/2024.05.06.592641

**Authors:** Arikta Biswas, Yuting Lou, Boon Heng Ng, Kosei Tomida, Sukhada Darpe, Zihao Wu, Thong Beng Lu, Isabelle Bonne, Chii Jou Chan

## Abstract

The maturation of functional eggs within the ovaries is essential for successful reproduction and organismal functions in mammals. Yet, despite its biological and clinical importance, the underlying mechanisms regulating folliculogenesis remain enigmatic. Here, we report a novel role of the surface-anchoring theca cells (TCs) in regulating follicle growth through mechanical signalling. Direct mechanical measurements reveal that these TCs are highly contractile and exert compressive stress to the follicular interior, potentially through active assembly of fibronectin scaffold around the follicles. Abolishing TC contractility disrupts fibronectin assembly, increases follicle size, and decreases intrafollicular pressure and viscosity. We further reveal that the granulosa cells (GCs) within the follicles exhibit spatial patterns of YAP signalling and proliferation, which appear to be decoupled. Transient manipulation of tissue pressure through bulk follicle compression, laser ablation or pharmacological perturbation of TC contractility leads to changes in GC YAP signalling, proliferation, and oocyte-GC communications, while long term abrogation of TC contractility leads to impaired follicle growth. Altogether, our study unveils the unique role of TC-mediated tissue pressure in ensuring robust mammalian ovarian folliculogenesis.

## Introduction

The maturation of functional oocytes within the ovarian follicles is undoubtedly one of the most significant developmental events in reproductive biology. The growth of follicles, or folliculogenesis, is essential for ensuring successful reproduction and regulating hormones for female sexual characteristics and early pregnancy^1–4^. Folliculogenesis begins with the primordial follicle where a single oocyte is surrounded by a layer of granulosa cells (GCs) and basement membrane (BM). Upon activation, they develop into primary follicles characterised by the formation of columnar GCs, which surround the oocyte with a glycoprotein shell of zona pellucida (ZP). The oocyte and the GCs maintain bi-directional communications through the transzonal projections (TZPs). The follicles then develop into secondary follicles with the formation of multi-layered GCs and an external layer of spindle-shaped theca cells (TCs). As the follicles grow, a large fluid-filled lumen forms within the GC layers which ultimately leads to follicle rupture and release of the oocyte, a process known as ovulation.

While past molecular genetics studies have identified genes that are critical for folliculogenesis^5–8^, the underlying mechanisms regulating follicle growth remain enigmatic. In recent years, new evidence has emerged showing that the ovary is a mechanically responsive organ^9^ and that mechanical signalling can impact follicle dynamics and development^10–12^. For example, it has been reported that the fragmentation of ovaries can disrupt the Hippo signalling pathway and promote follicle growth^13–15^, while mechanical stress imposed by the extracellular matrix (ECM) play a role in regulating primordial follicle dormancy^16,17^. Changes in ECM stiffness during ovarian ageing has also been implicated in impaired oocyte quality^18^ and anovulation^19^. Other evidence come from *ex vivo* studies showing that the growth of isolated follicles is highly sensitive to the surrounding matrix stiffness^20^. However, despite these evidence, fundamental questions such as how mechanical forces are generated and transmitted within the follicles, and how these mechanical signals orchestrate morphogenesis and oocyte maturation, remain unclear.

A recent study revealed that the intra-follicular environment is characterised by distinct mechanical properties of TCs and GCs^21^. The TCs are indispensable for folliculogenesis as they are involved in the production of steroid hormones for ovulation^4,22–25^. Abnormalities in TC steroid secretion can lead to polycystic ovary syndrome^26–28^, a prominent cause of female infertility^29,30^, and hyperthecosis^31,32^, a condition usually affecting postmenopausal women and causing virilization^33^. TCs have also been implicated in early menopause in reproductive-aged women^34,35^. Yet, apart from hormonal regulation, the structural and mechanical functions of TCs remain largely unknown. In this study, we investigated the mechanical interactions between TCs and granulosa cells during secondary follicle development. Using *in vitro* and *ex vivo* approaches, combined with quantitative imaging, biophysical tools, and molecular perturbations, we revealed the novel roles of contractile theca cells in exerting active compressive stress to tune tissue pressure and mechanics, thereby regulating somatic cell signalling and follicle growth.

## Results

### Ovarian theca cells are highly contractile

The surface cells in spherical tissues such as embryos and cell aggregates are often found to be highly stretched and contractile^36,37^. We thus hypothesised that the peripheral spindle-shaped TCs may exert strong contractile forces around the ovarian follicles. We immuno-stained ovarian tissue slices and isolated secondary follicles targeting phosphorylated myosin light chain 2 (pMLC), which has been reported to be a good proxy for actomyosin contractility^36^. We found that the TCs express high amounts of pMLC compared to the minimal levels observed in the GCs, both *in situ* (Figures 1A-B) and *ex vivo* (Figures 1C-D), suggesting that the TCs are indeed contractile. Inhibition of actomyosin contractility with blebbistatin (Blebb) led to a decrease in TCs’ pMLC expression while hyperactivation of contractility with lysophosphatidic acid (LPA) did not increase the TCs’ pMLC expression further, both *in situ* (Figures S1A-B) and *ex vivo* (Figures S1C-D). We also observed an increase in the pMLC expression at TCs with increased follicle size (Figures 1B, right and 1D, right), suggesting that the TC layers become more contractile as the secondary follicles develop. Though the oocyte cortex expressed some levels of pMLC expressions, its intensity did not change upon actomyosin perturbations (Figures S1D-E), suggesting that the impact of the drugs is mainly specific to the outer contractile TCs.

**Figure 1:**
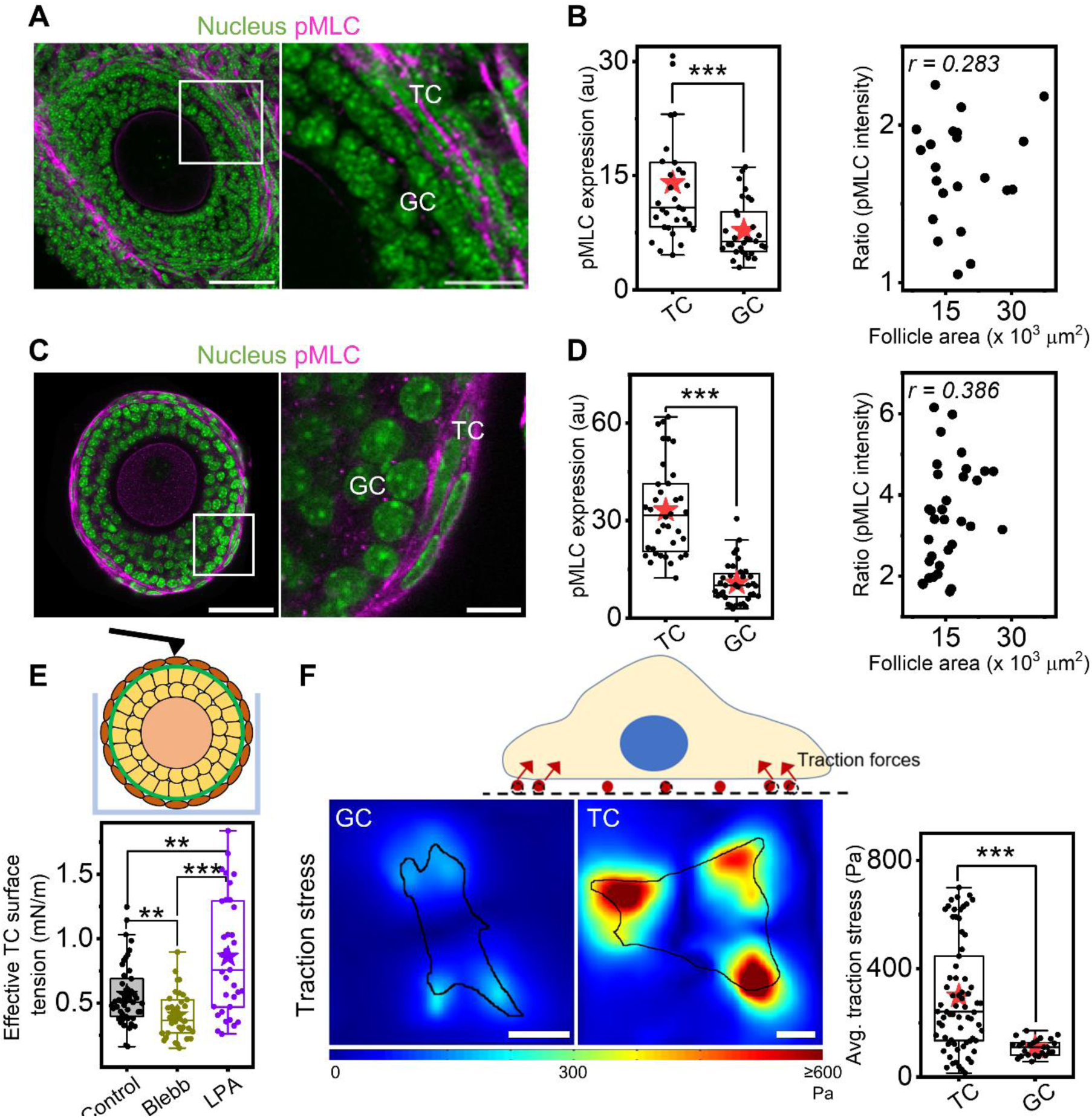
Ovarian theca cells are highly contractile. A) Left: Representative image of an ovarian tissue slice labelled with DAPI (nucleus, green) and immuno-stained with phosphorylated-myosin light chain (pMLC, magenta). Scale bar: 50 µm. Right: inset shows the zoomed-in region marked in white. Scale bar: 20 µm. B) Left: Boxplots of pMLC intensities in TCs and GCs *in situ*. Right: Scatter plot of pMLC intensity ratios as a function of follicle size. N = 2, n = 31 follicles. C) Left: Representative image of an isolated secondary follicle labelled with DAPI (green) and immuno-stained with pMLC (magenta). Scale bar: 50 µm. Right: inset shows the zoomed-in region marked in white. Scale bar: 10 µm. D) Left: Boxplots of pMLC intensities in TCs and GCs *ex vivo*. Right: Scatter plot of pMLC intensity ratios as a function of follicle size. N = 3, n = 36 follicles. E) Top: Schematic of AFM-based indentation on a follicle in a microwell to measure TC surface tension. Bottom: Boxplots of effective TC surface tension in control, Blebb, and LPA-treated samples. N = 3, n = 45 (control), 35 (Blebb, LPA) follicles. F) Top: Schematic of traction force microscopy. Bottom left: Representative traction stress maps for isolated GCs and TCs *in vitro*. Outline of the cells are marked in black. Scale bar: 10 µm. Bottom right: Boxplot of average traction stress (per cluster) for TCs and GCs. N = 25 cells, n = 78 clusters. Significance was determined by Mann-Whitney U test. ** p < 0.01; *** p < 0.001.

We next measured the surface tension of TCs in the secondary follicles using atomic force microscopy (AFM, Supplementary Methods). We found that the TCs exhibit an effective surface tension of 0.51±0.20 mN/m (Figure 1E), similar to that found for stretched cells in living tissues^37^. Follicles treated with Blebb and LPA showed a significant decrease and increase in the measured TC surface tension, respectively (Figure 1E). This is consistent with cell rounding or stretching associated with surface tension release or increase (Figures S1F-G). We then isolated primary TCs from bulk ovaries^38,39^ to check if the high contractility is an intrinsic feature of the TCs. Consistent with previous reports which show alkaline phosphatase (ALP) positive staining on the TCs in pre-ovulatory follicles^40–42^, we noted ALP expression at the periphery of follicles in ovarian slices (Figure S2A). Isolated TCs expressed more ALP and appeared more elongated and spread out on 2D substrates compared to the smaller and more cuboidal GCs that are ALP-negative (Figures S2B-D). Using traction force microscopy, we found that the TCs exert significantly higher traction stresses than the GCs (Figure 1F) which is correlated with their spread area (Figure S2E), indicating they are indeed intrinsically more contractile than the GCs.

Altogether, based on our findings *in situ*, *ex vivo* and *in vitro*, we conclude that the TCs are highly contractile, which can potentially regulate follicle development through mechanical signalling.

### Theca cells exert compressive stress to module follicle mechanics and pressure

We hypothesized that the contractile TCs may exert compressive stress to regulate follicle size and functions. To directly measure the compressive stress imposed by the TCs, we allowed secondary follicles to attach to deformable gelatin beads and tracked the bead-follicle pairs for two days by time-lapse imaging. We observed that the TCs migrated from the follicles to the beads and uniformly enwrapped the beads within 12 hours. Using dextran-based osmotic compression assay^43^, we found that the beads have an average bulk modulus (a measure of compressibility) of 19.4±6.3 kPa (Figure 2A, Methods). This information, combined with the tracking of changes in bead volume during TC enwrapment (Figure 2B, left), allowed us to uniquely determine the compressive stress exerted by the TCs on the beads to be ∼2 kPa. The compressive stress decreased or increased significantly with Blebb or LPA treatment (Figure 2B, right), respectively, suggesting that the compressive stress originated from TC contractility.

**Figure 2:**
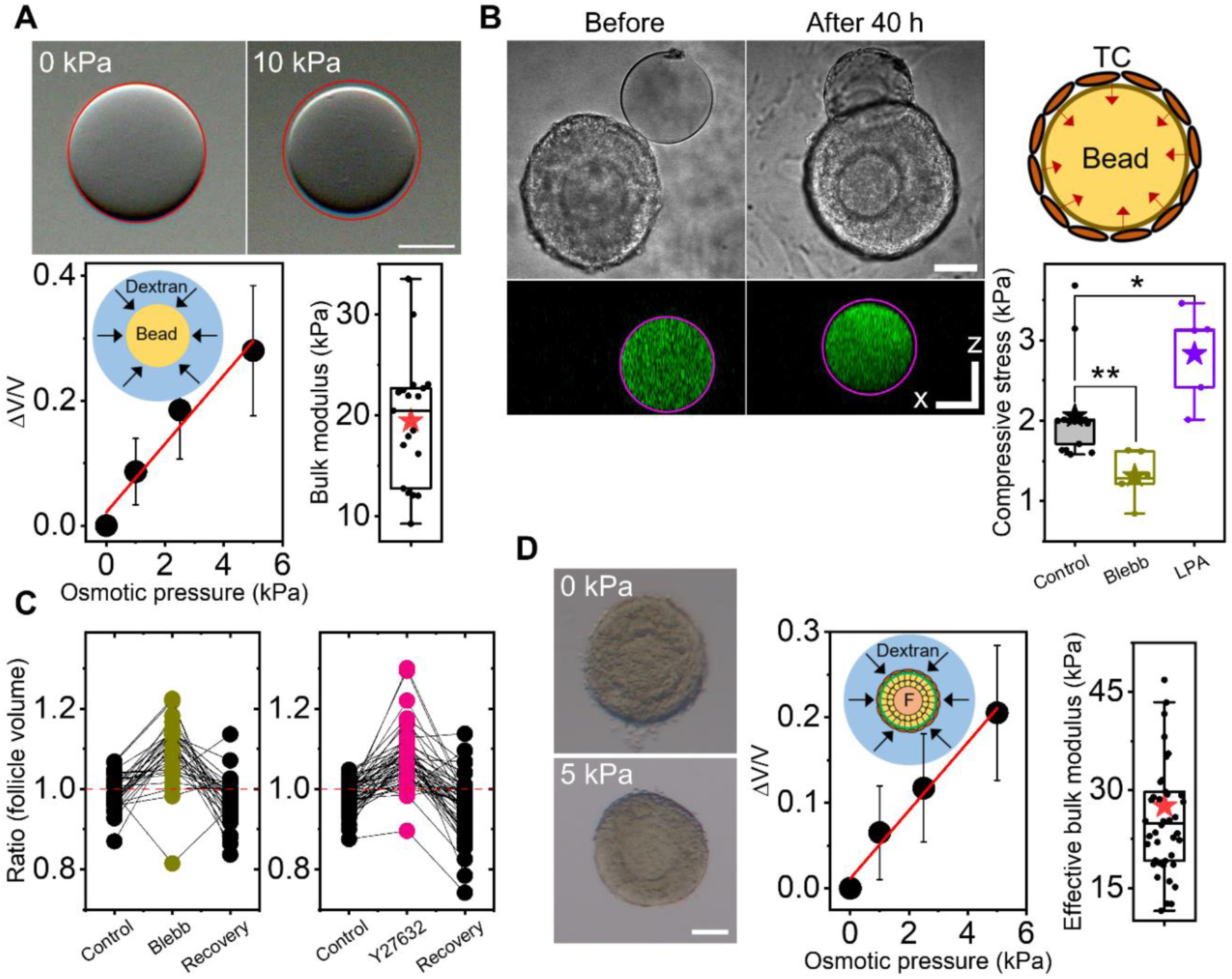
Theca cells generate compressive stress to regulate follicle size. a) Top: Representative images of a bead under different osmotic stress. The outline of the initial bead boundary is marked in red. Scale bar: 50 µm. Bottom left: Plot of average relative change of bead volume against osmotic stress (black symbols) and the linear fit (red line). Bottom right: Boxplot of measured bulk modulus of beads. N = 2, n = 20 beads. B) Left: Representative images of a bead before and after TC enwrapment – brightfield (top) and orthogonal view (bottom). The outline of the initial bead boundary is marked in magenta. Scale bar: 50 µm. Right: Boxplots of compressive stress measured in control, Blebb, and LPA-treated TCs. N > 2, n = 6-17 bead-follicle pairs. C) Boxplots of follicle volume change upon perturbations of contractility (left: Blebb; right: Y27632) and washout (recovery). N = 3, n = 41 (Blebb), 57 (Y27632) follicles. D) Left: Representative images of a secondary follicle under different osmotic pressure. Scale bar: 50 µm. Mid: Plot of average relative change in follicle volume against osmotic stress (black symbols) and the linear fit (red line). Right: Boxplot of measured bulk modulus of secondary follicles. N = 2, n = 42 follicles. Error bars in A) and D) represent standard deviation. Significance was determined by Mann-Whitney U test in (B). * p < 0.05. ** p < 0.01.

We next considered if perturbing TC contractility directly impacts follicle size. On tracking every follicle, we observed that transient inhibition of contractility with Blebb (Figure 2C, left) or Y27632 (Figure 2C, right) for 30 mins led to an increase in follicle volume. A wash out of both inhibitors over similar timescales led to a restoration of follicle volume, suggesting that the volume regulation by TC-mediated contractility is fast, global and reversible. Using a similar dextran-based compression assay, we determined the effective bulk modulus of secondary follicles to be ∼27.5±13.4 kPa (Figure 2D). This, together with the measured volumetric strain of ∼0.1 upon perturbation (Figure 2C), allowed us to infer the compressive stress exerted by TCs on secondary follicles to be ∼2.75 kPa, consistent with that measured using the bead-follicle assay.

To investigate if changes in follicle volume by TC-mediated compressive stress affect intrafollicular pressure and bulk mechanics, we performed AFM indentations on secondary follicles under various perturbations, using large beads (Figures 3A). We found that while LPA treatment did not change tissue elasticity (Figure 3B and Supplementary Information) and effective pressure (Figure 3D) as compared to that of the controls, blebbistatin treatment led to a significant decrease in both parameters. Furthermore, we observed a large hysteresis between the approach and retraction curves in Blebb-treated follicles (Figure 3C), indicating that the release of compressive stress from TC relaxation leads to increased stress dissipation and a more fluid-like state. To confirm the change in follicle viscosity, we extracted the tissue viscosity by fitting the AFM data to a Maxwellian viscoelasticity model (Star Methods and Supplementary Information). Our results indeed revealed a decrease in follicle viscosity upon Blebb treatment (Figure S3A).

**Figure 3:**
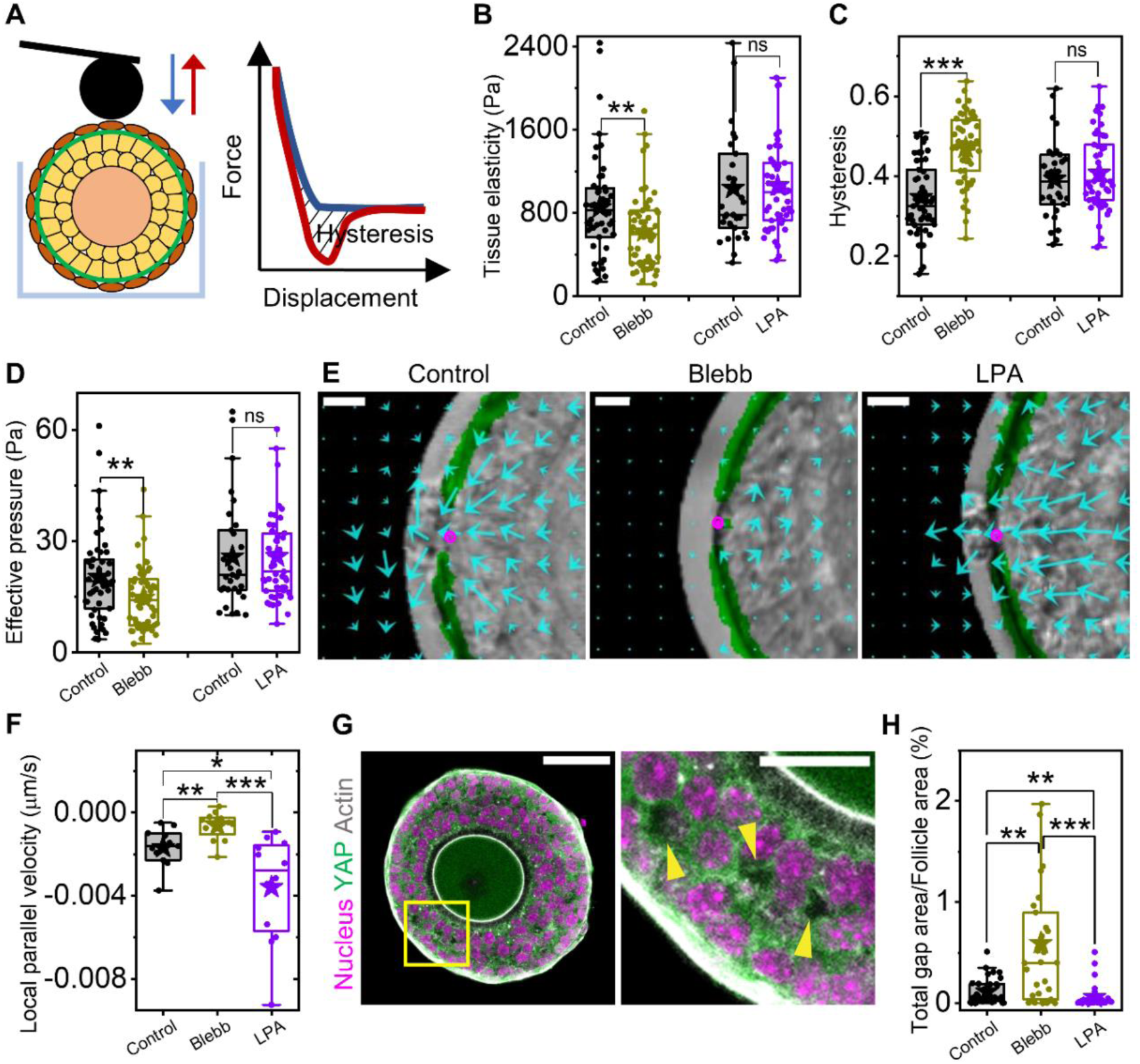
Compressive stress modulates follicle mechanical properties. (A-D) Schematic of AFM approach to measure follicle mechanical properties from the approach (blue) and retraction curve (red) (A), showing how this yields the effective follicle elasticity (B), hysteresis (C), and effective pressure (D) in various conditions. N = 5, n = 51 (control), 55 (Blebb); N = 2, n = 31 (control), 51 (LPA) follicles. E) Zoomed-in representative displacement vector maps overlaid on CNA35 (green) marked follicles (brightfield) in control, Blebb, and LPA treatments; ablation site is marked by magenta circles. Scale bar: 20 µm. F) Boxplots of local parallel velocity in the three conditions. N = 4, n = 11-12 follicles each. G) Left: Representative image of an isolated follicle labelled with DAPI (nucleus, magenta), Phalloidin (actin, grey), and immuno-stained with YAP (green). Scale bar: 50 µm. Right: Zoomed-in image of the yellow box marked on the left. Yellow arrowheads demarcate interstitial gaps. Scale bar: 20 µm. H) Boxplots of the ratios of total interstitial gaps to follicle area for follicles in various conditions. N = 3, n = 34 (control), 31 (Blebb), 32 (LPA) follicles. Significance was determined by Mann-Whitney U test. ns: p > 0.05; * p < 0.05. ** p < 0.01; *** p < 0.001

To further validate the role of TC contractility on intrafollicular pressure, we performed two-photon laser ablations by making a point cut at the follicle periphery, followed by tracking of tissue outflow. Following ablation, we observed a rapid displacement of GCs towards the ablation site (Figure 3E). By quantifying the GC flow near the cut region (local parallel velocity, Methods) in various conditions, we observed a significant attenuation of GC flow upon Blebb treatment while LPA led to increased GC outflow compared to the controls (Figure 3F). These data confirm that Blebb or LPA treatment led to reduced or increased intrafollicular pressure, respectively. Finally, we found an increase in total interstitial gap area within the Blebb-treated follicles (Figures 3G-H and S3B), suggesting that reduced tissue packing may lead to the overall reduction in tissue elasticity and pressure while enhancing tissue fluidity. By contrast, LPA led to only a slight further decrease in interstitial gap area, suggesting that the follicles in their native state are already tightly packed and are not susceptible to further compression, consistent with the lack of a change in follicle volume (Figure S3C) and mechanics (Figure 3B-D).

### Theca cell contractility mediates fibronectin-scaffold assembly

To gain structural insights on how forces are transmitted by the TCs, we immuno-stained for adherens junctional proteins such as N-Cad and E-Cad (Figure 4A). In contrast to the GCs which expressed these junctional markers, the TCs are devoid of these proteins, suggesting that they are less epithelial in nature and resemble more mesenchymal and fibroblast-like cells. Indeed, the theca layers expressed abundant fibronectin (FN) compared to the GCs (Figures 4B-C), and their expressions appeared to increase with follicle development (Figure 4C, right). This pattern was preserved in isolated follicles as well (Figures 4D-E). Interestingly, while the BM has been reported to be enriched with FN in past studies^44^, we found the FN layer to be physically separated from the collagen matrix at the BM, as shown by localisation studies (Figure 4F). In addition, ultrastructural studies using Scanning Electron Microscopy (SEM) revealed a distinct matrix-like layer separating the basal TCs from the BM (Figure 4G and S4A). Since FN is not expressed in the primordial and primary follicle stages when the TCs are absent^44,45^, we hypothesized that the TCs might be actively secreting the FN which could be a constituent of the matrix between the basal TCs and BM. Though the average thickness of the matrix (456±152 nm) did not change with increased follicle size, we observed an increase in the variability of the matrix thickness with follicle development (Figure S4A-C).

**Figure 4:**
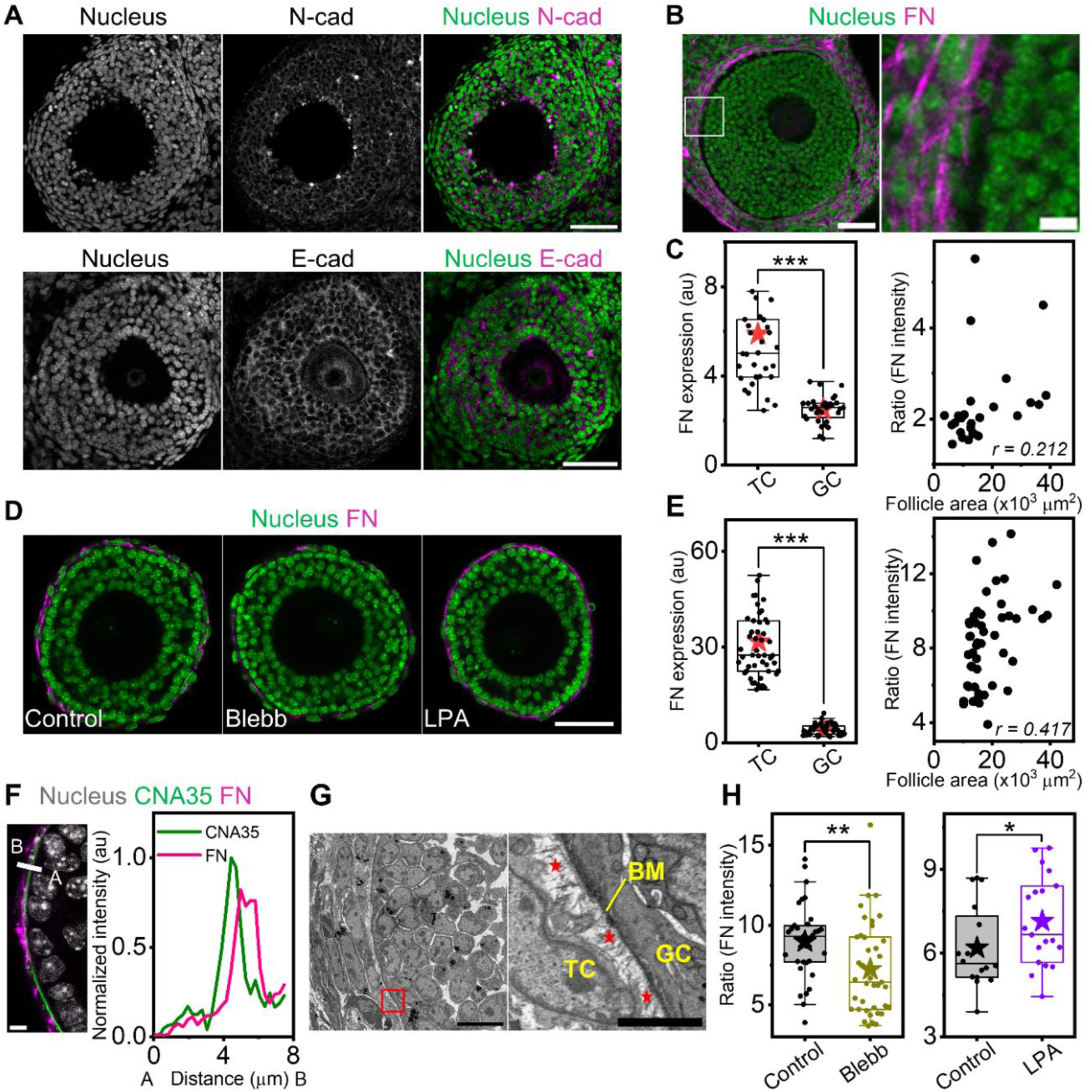
Theca cell contractility regulates fibronectin scaffold formation. A) Representative images of ovarian slices immuno-stained with N-cad (top) or E-cad (bottom) junctions. Scale bar: 50 µm. B) Left: Representative image of an ovarian slice labelled with DAPI (green) and immuno-stained with FN (magenta). Scale bar: 50 µm. Right: Zoomed-in region of the marked white box. Scale bar: 5 µm. C) Left: Boxplots of FN intensity in TCs and GCs *in situ*. Right: Plot of TC FN expression against follicle size. N = 2, n = 32 follicles. D) Representative images of isolated secondary follicles labelled with DAPI (green) and immuno-stained with FN (magenta) in various conditions. Scale bar: 50 µm. E) Left: Boxplots of FN intensity in TCs and GCs within control follicles *ex vivo*. Right: Plot of TC FN expression against follicle size. N = 4, n = 50 follicles. F) Left: Zoomed-in section of a follicle 18mmune-stained with FN (magenta) and stained with DAPI (grey) and CNA35 (green). Scale bar: 10 µm. Right: Plot of intensity profile for the line scan marked in white (left image) shows a physical separation of FN and collagen at the BM site. G) Left: Representative SEM image of a section of a follicle. Right: Zoomed-in section of the red box marked on left. Red asterisks indicate the fibronectin-rich matrix between the BM and the basal TCs. Scale bars: 10 and 2 µm respectively. H) Boxplots of normalised FN intensity of TCs under Blebb (left) and LPA (right) treatments. N = 3; n = 33 (control), 39 (Blebb). N = 2; 16 (control), 20 (LPA) follicles. Significance was determined by Mann-Whitney U test. * p < 0.05; ** p < 0.01; *** p < 0.001.

Following a recent finding that contractile cancer-associated fibroblasts (CAFs) can produce fibronectin scaffolds around tumour cells for force transmission^46^, we were curious if perturbing TC contractility affects fibronectin expression in ovarian follicles. We observed that blebbistatin treatment led to a significant reduction of FN expression at the TC layer *ex vivo* and *in situ*, while LPA treatment led to increased FN expression *ex vivo* but not *in situ* (Figure 4H, S4D-E). Here, the short timescale (4 hr) for FN remodelling in response to pharmacological perturbations may be due to the resemblance of follicular fibronectin to those found in fetal development rather than that of the adult tissues^47^. Conversely, treatment of follicles with RGDS peptides to inhibit TC-adhesion to fibronectin did not impact FN or pMLC expressions at the TCs (Figure S4F-H). Together, these data indicate that though actomyosin perturbations could impact fibronectin scaffold around the follicles, TC-fibronectin coupling is not essential for maintaining TC contractility and FN integrity.

### Granulosa cells show spatial patterns of proliferation and YAP signalling

We next investigated if the signalling landscape within the follicles are sensitive to the mechanical environment. Inspired by studies showing that cell proliferation could be tuned by mechanical stress in cancer spheroids^48^, we immuno-stained tissue slices and isolated follicles with Ki67, a known cell proliferation marker. We observed that the GCs in contact with the BM (basal GCs) were significantly less proliferative than the GCs surrounding the oocyte (oocyte GCs) in both tissues (Figures 5A-B). A similar pattern of differential proliferation between the basal- and oocyte GCs was also observed when follicles were labelled with EdU, another proliferation marker (Figures S5A-B).

**Figure 5:**
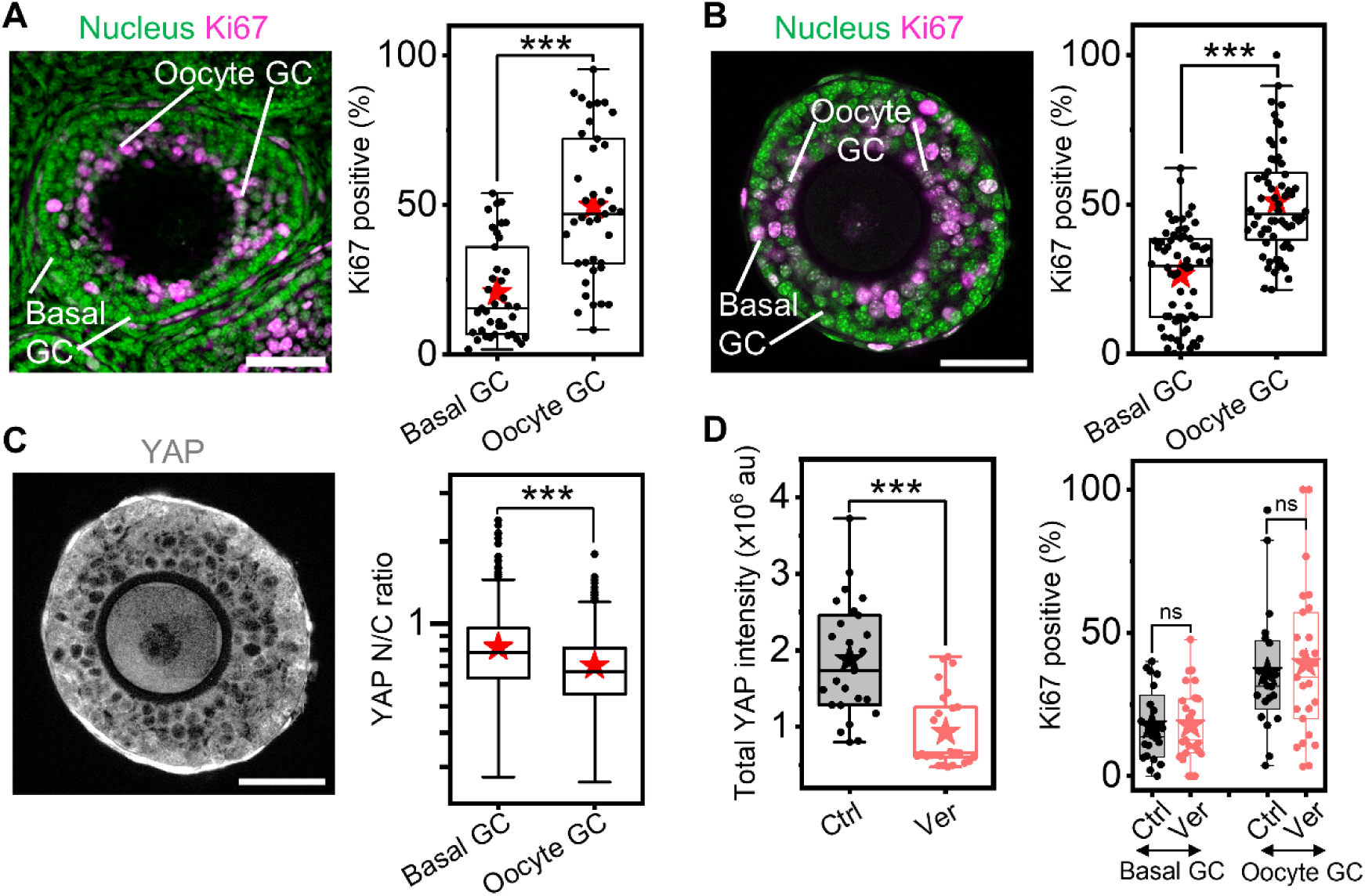
Granulosa cells show differential proliferation and YAP signalling within the follicle. A) Left: Representative image showing an ovarian slice labelled with DAPI (green) and 20mmune-stained with Ki67 (magenta). Scale bar: 50 µm. Right: Boxplots for percentage of Ki67^+^ basal and oocyte GCs within secondary follicles *in situ*. N = 3, n = 39 follicles. B) Left: Representative image showing an isolated secondary follicle stained with DAPI (green) and Ki67 (magenta). Right: Boxplots for percentage of Ki67^+^ basal and oocyte GCs *ex vivo*. C) Left: Representative image of the same follicle in (B) immuno-stained with YAP. Right: Boxplots of YAP N/C ratios (log scale) in basal and oocyte GCs *ex vivo*. N = 4, n = 67 follicles. Scale bar: 50 µm. D) Left: Boxplots of total YAP intensity in control and verteporfin-treated isolated follicles. Right: Boxplots for percentage of Ki67^+^ basal and oocyte GCs *ex vivo* in control and verteporfin conditions. N = 2, n = 25 (control), 27 (verteporfin) follicles. Significance was determined by Mann-Whitney U test. ns: p > 0.05; *** p < 0.001.

We also immuno-stained the secondary follicles with YAP, a transcriptional co-activator that is known to be mechanosensitive^49^ and is important for ovarian folliculogenesis^14^. We found that the YAP nuclear-to-cytoplasmic (N/C) ratio for basal GCs was significantly higher than that of the oocyte GCs (Figure 5C), raising the intriguing possibility of the presence of a mechanical stress gradient within the follicle. Of note, we observed an anti-correlation between YAP signalling and cell proliferation. To investigate this, we treated follicles with verteporfin, which is known to inhibit YAP nuclear translocation and reduce cell proliferation^50^. While verteporfin led to an overall decrease in YAP expression for the GCs in the follicles (Figure 5D, left), the differential pattern for Ki67 signalling remained unchanged (Figure 5D, right). This suggests that the Hippo signalling pathway may not dictate GC proliferation during ovarian follicle development.

### Transient mechanical stress impacts GC signalling and oocyte-GC communications

Next, we investigated if perturbing compressive stress pressure via changing osmotic pressure, TC contractility or BM stiffness could influence the proliferation and YAP signalling patterns of the oocyte- and basal GCs. Focusing on the Ki67 signals, we observed that the proliferation potential of basal GCs did not change upon all perturbations over a short timescale (∼30 mins) (Figures 6A-B). Transient increase in compressive stress via osmotic pressure (10 kPa) or hypercontractility of TCs (LPA) led to a striking reduction of Ki67^+^ cells among the oocyte GCs (Figures 6A-B). However, a transient release of compressive stress (tissue pressure) either by contractility inhibition with Blebb or Y27632, or by BM degradation with collagenase (Figure S6A) showed no impact on the proliferation of oocyte GCs. Using EdU pulse-chase assays, we saw that the effect of the perturbations impacted the basal GCs more than the cells towards the core. Increased mechanical stress over short timescale (chased for 30 mins) reduced EdU proliferation as well (Figures S5A-B).

**Figure 6:**
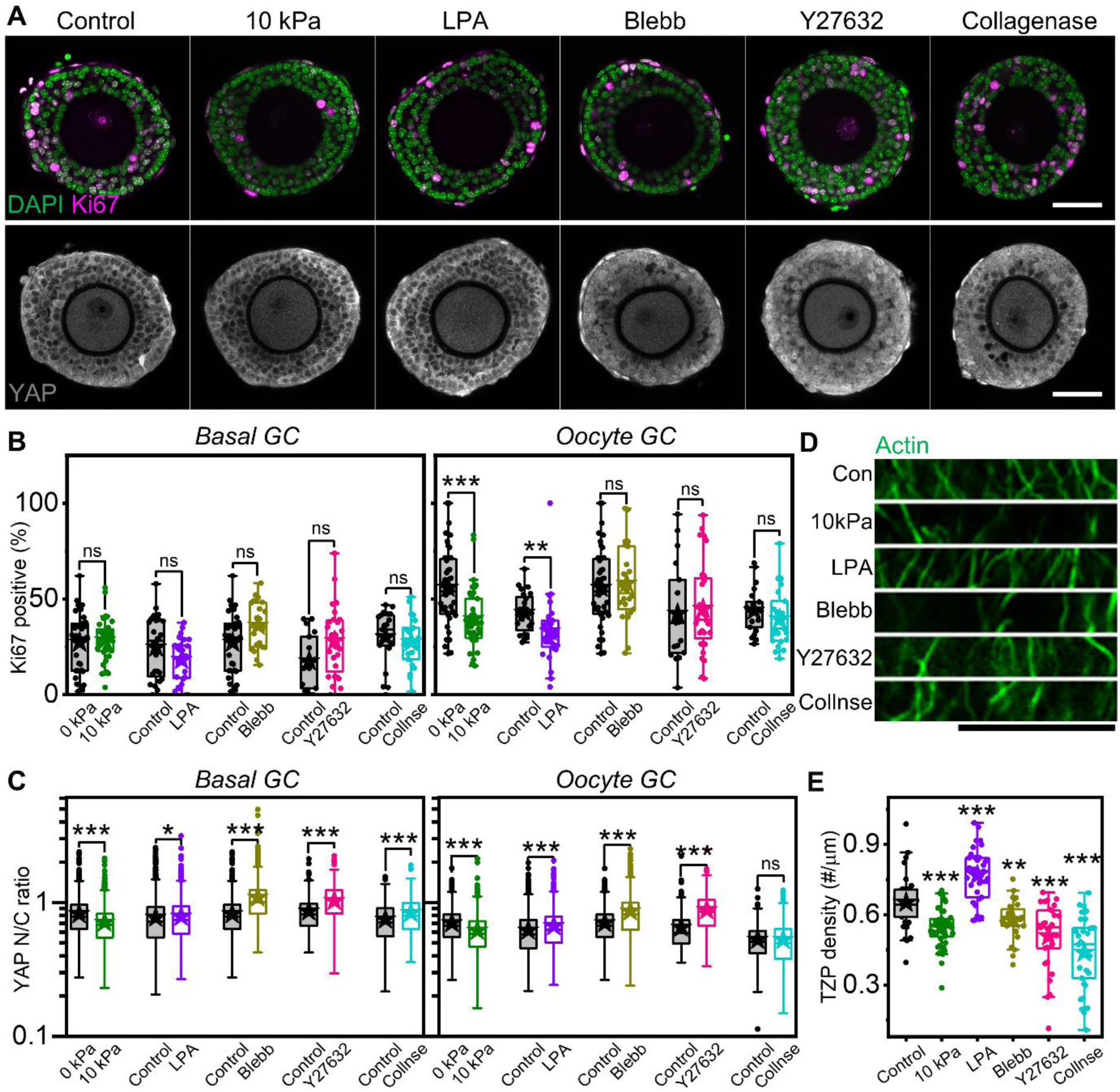
Transient perturbation of mechanical stress impacts granulosa cell signalling and oocyte GC communications. A) Representative images of follicles stained with DAPI and immunolabelled with Ki67 (top), and immunolabelled with YAP (bottom) upon transient perturbation of mechanical stress. Scale bar: 50 µm. B) Boxplots for percentage of Ki67^+^ basal and oocyte GCs in various perturbations. C) Boxplots of YAP N/C ratios (log scale) in basal and oocyte GCs in various perturbations. N =3, n = 28 (control), 38 (10 kPa), 29 (LPA), 26 (Blebb), 38 (Y27632), 23 (Collnse) follicles. D) Representative images of actin transzonal projections between the oocyte and oocyte GCs for follicles in various perturbations, with corresponding boxplots of TZP number density shown in (E). Scale bar: 10 µm. N = 3, n = 25 (control), 48 (10 kPa), 30 (LPA), 28 (Blebb), 35 (Y27632), 32 (Collnse) follicles. Significance was determined by Mann-Whitney U test. ns: p > 0.05; * p < 0.05; ** p < 0.01; *** p < 0.001.

Transient mechanical perturbations have a more striking impact on YAP signalling of GCs (Figures 6A and 6C). Osmotic compression (10 kPa) led to an increase in YAP cytoplasmic localization in both basal- and oocyte GCs. However, enhancement of TC contractility with LPA treatment appeared to increase the YAP N/C ratios. Stress relaxation by Blebb, Y27632 or collagenase perturbations led to a significant increase in YAP nuclear translocation. Though we observed no change in the YAP localization of GCs at the core of the follicles on transient collagenase treatment, the impact on YAP nuclear translocation of both basal- and oocyte GCs was more pronounced over longer treatment of 2 hours, although proliferation was not affected (Figure S6B). pMLC expression at the TCs reduced upon BM disruption, indicating a loss of TC contractility in collagenase-treated follicles (Figure S6C). Next, we released the tissue pressure by laser ablation at the BM and found that the YAP N/C ratios of GCs in these follicles were higher compared to the controls (Figures S6D-E). Altogether, our results support that the modulation of intrafollicular pressure by physical perturbations, BM degradation or altered TC contractility can all individually regulate intra-follicular Hippo signalling pathway at short timescales (Figure S6F).

We then focussed on the impact of transient mechanical perturbations on transzonal projections, which are filopodia-like structures connecting the oocyte GCs to the oocyte that are essential for its growth^51^. We found that the number density of TZPs reduced significantly upon osmotic compression, contractility inhibition and BM disruption (Figures 6D-E), which is not correlated with the minimal change in the thickness of zona pellucida in these conditions (Figure S5C). We also examined how the oocyte volume changes upon various perturbations and observed that while osmotic compression led to a significant decrease in oocyte volume (∼10%), there was minimal impact of contractility perturbations and BM disruption on the oocyte volume (Figure S5D).

### Mechanical stress is required for follicle growth

To examine the functional consequence of TC contractility on follicle growth, we cultured follicles within 3D alginate hydrogels for up to three days under various pharmacological perturbations. By day 3, we observed that the average diameter of follicles under LPA treatment was higher than that of the controls, particularly for follicles with initial size of less than 150 µm (Figure 7A). While a small dosage of blebbistatin (5 µM) had no impact on follicle growth (Figure S7A-C), a higher dosage of blebbistatin (20 µM) led to impaired follicle growth by day 3, particularly for follicles with initial size larger than 150 µm (Figure 7A). A similar, albeit less pronounced effect was seen in follicles treated with Y27632. Compared to the controls where typically ∼15% of the follicles showed follicle rupture and oocyte extrusion during culture, such events were more frequently observed with LPA treatment (∼30%) but less with Blebb and Y-27632 treatment (< 5%) (Figure 7B) This suggests that follicle ruptures could be a consequence of enhanced tissue pressure, as observed from the laser ablation studies (Figure 3F).

**Figure 7:**
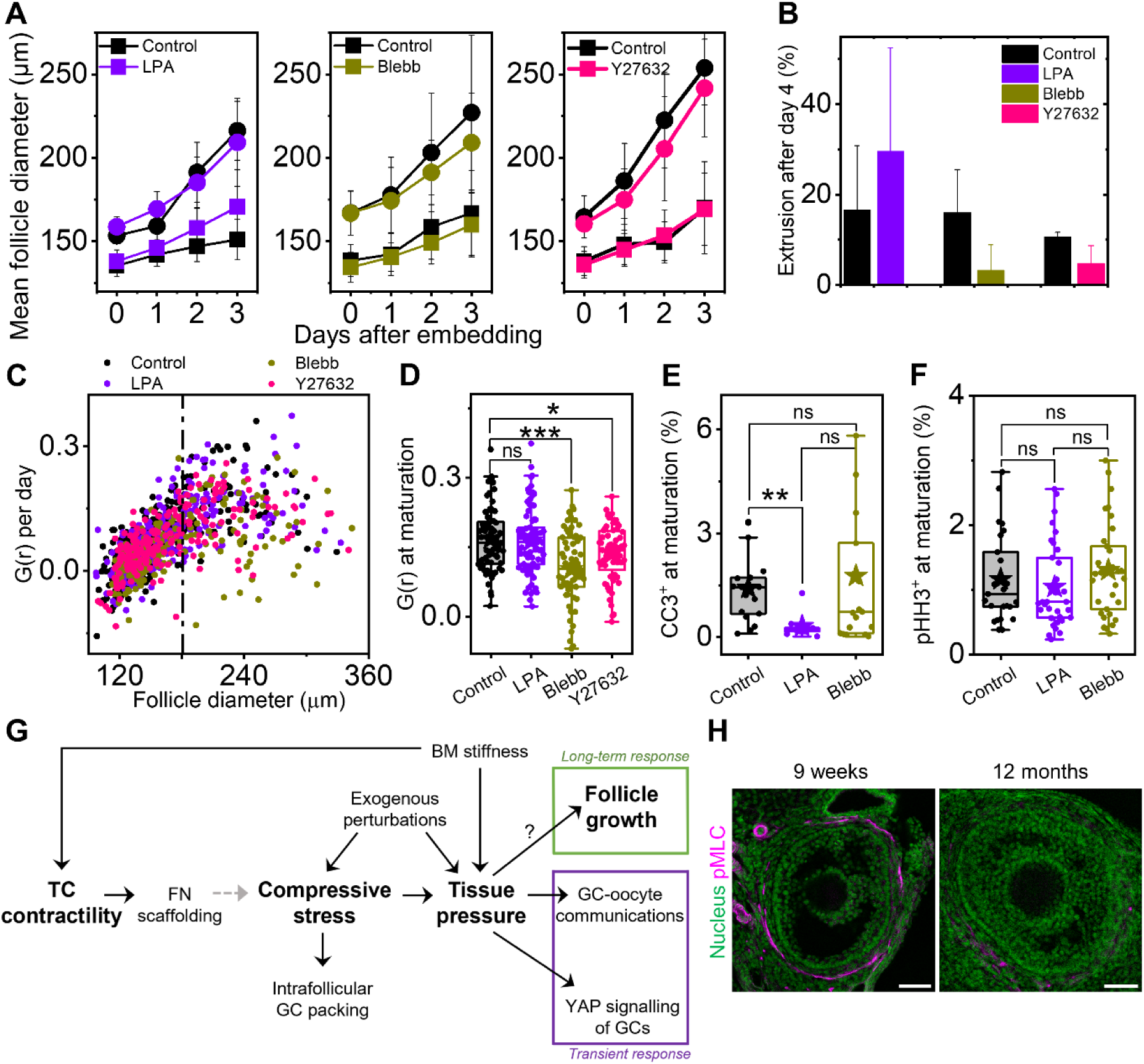
Attenuation of TC contractility leads to reduced follicle growth. A) Plot of follicle diameters in controls and various contractility perturbations. N = 3-5, n = 172 (control), 86 (LPA), 89 (Blebb), 81 (Y27632) follicles. Error bars represent standard deviation. Square and circle symbols represent follicles with diameters smaller and larger than 150 µm respectively. B) Percentage of extrusion events upon contractility perturbation. Bars represent mean of ruptures in an experiment and error bars represent standard deviation. C) Scatter plot of growth rates per day as a function of follicle size in different conditions. Dashed line marks the transition to maturation phase. D) Boxplots of maturation growth rates for follicles in different conditions. n = 72 (control), 86 (LPA), 65 (Blebb), 64 (Y27632) follicles. E) Boxplots of CCL3^+^ GCs in matured follicles cultured in different conditions. N = 2, n = 25 (control), 28 (LPA), 32 (Blebb) follicles. F) Boxplots of pHH3^+^ GCs in matured follicles after culture in different conditions. N = 2, n = 38 (control), 43 (LPA), 42 (Blebb) follicles. G) Schematic showing changes in follicle pressure through extrinsic mechanical perturbations or contractile TCs can impact GC mechanical signalling, oocyte-GC communications, and long-term follicle growth. H) Representative images of follicles immune-stained with DAPI (nucleus, green) and pMLC (magenta) in young (9 weeks) and aged (12 months) ovaries. Significance was determined by Mann-Whitney U test. ns: p > 0.05; * p < 0.05; *** p < 0.001.

As the follicle growth kinetics appeared to depend on the initial size (Figure 7A), we sought to develop an integrative approach to combine the growth kinetics of all follicles of arbitrary sizes into a master curve, thus allowing quantitative comparison of follicle growth in various conditions. By plotting the follicle growth rate per day versus its size (Figure 7C), we found that the follicle growth is characterised by two phases: an initial pre-maturation phase where the growth rate increased linearly with its size, followed by the maturation phase (*D* > 180 µm) where the grow rates reach a terminal value. While LPA-treated follicles showed no difference in their maturation growth rates from the controls, follicles treated with contractility inhibitors showed attenuated growth at the maturation phase (Figure 7D), further confirming that reduced compressive stress leads to impaired follicle growth.

Reduced growth rates of follicles may be attributed to a decrease in cell proliferation or increased apoptosis. To this end, we stained post-cultured follicles for cleaved caspase3 (CC3), an indicator for apoptosis, and phospho-histone H3 (pHH3), a marker for mitosis. Intriguingly, we observed no difference in the number of apoptotic or mitotic GCs in control follicles and Blebb- and LPA-treated samples (Figures 7E-F and S7D), though cell death was significantly reduced when cultured in LPA. We also found no difference in the number of GCs at telophase in the three conditions (Figures S7E), suggesting that Blebb treatment did not cause deleterious effects such as delayed cytokinesis. Overall, our data revealed that an optimal amount of TC-mediated compressive stress is required for 3D follicle growth (Figure 7G).

## DISCUSSION

In the past decades, we have made significant progress in understanding the roles of oocyte-granulosa cell signalling pathways^52,53^, ECM and stroma^54,55^ in ovarian biology. However, the origin and functions of the theca cells that make up the periphery of preantral follicles remain poorly understood. While it has long been proposed that the theca externa may exert contractile forces to aid ovulation^56,57^, existing studies on TCs remain largely limited to their steroidogenic functions^4,22–25^ with little examination on their potential mechanical roles. This is highly pertinent given the recent evidence that mechanical cues in the follicle microenvironment can regulate diverse follicle functions, from activation to growth and ovulation^12,14,16,20^. In this study, using a combination of biophysical, bioengineering, and molecular approaches, we characterised the detailed TC mechanics and unravelled its integral role in exerting compressive stress to regulate intrafollicular pressure, granulosa cell signalling and follicle growth (Figure 7G).

We found that the TCs of murine secondary follicles are highly contractile, with values of surface tension similar to those reported for highly stretched surface cells in living tissues^37^. The intrinsic contractile nature of TCs, in contrast to the GCs, corroborates with recent optical elastography study showing that the TC shell possesses distinct mechanical stiffness compared to the GCs^21^, potentially acting as a mechanical cage to protect the oocyte from excessive deformation. We found that the TCs appear to be fibroblast-like cells that are devoid of adherens junction proteins and are capable of secreting FN networks in a contractility-dependent manner. This resembles a recent finding that contractile CAFs can form a fibronectin-rich capsule around tumour cells and exert compressive stress to trigger mechanotransduction in tumors^46^. The increase in the variability of FN-matrix thickness (Figure S4C) could be due to the differentiation of basal TCs into theca interna that undergoes vascularisation^22,58–61^. Interestingly, FN assembly has been shown to be mechanosensitive to tissue strain, as in the case of blastocoel expansion during early *Xenopus* development^62^. Whether follicle growth in turn generates tissue strain on TCs to trigger mechanosensing and FN assembly is unclear and constitutes an exciting topic for future research.

We demonstrate, using multiple biophysical tools, that transient abolishment of TC contractility leads to increased follicle size and concomitant decrease in tissue elasticity, viscosity, and effective pressure, which correlate with less frequent follicle rupture events in 3D culture. The effect is less pronounced in the case of hyperactivation of TC contractility using LPA, suggesting that follicles in their native state are close to a maximally compact state that render them less susceptible to further compression. We also report, for the first time, the presence of spatial patterns of YAP signalling and proliferation within ovarian follicles (Figure 5). Contrary to what has been reported in *in vitro* studies^63^, we observed an anti-correlation between YAP signalling and Ki67, which indicates that Ki67 signalling does not act downstream of YAP in ovarian follicles. The origin of the spatial patterns in GC signalling is unclear, and we propose that this could be due to direct biochemical or mechanical signalling from the BM or oocyte, or the presence of a mechanical stress gradient within the follicles. Cell shape control may be another factor, given that the basal GCs appear highly packed and columnar while the oocyte GCs appear spherical.

While abolishing TC contractility did not impact the GC proliferation as much, these perturbations significantly increased nuclear YAP localization of GCs, potentially due to the disruption of GC contacts with reduced tissue pressure and follicle swelling^64–66^. While we cannot rule out the direct impact of contractility perturbations on GC YAP signalling, our findings that both laser ablation and BM degradation (collagenase) lead to enhanced YAP nuclear transport of the GCs at short timescales demonstrate that GCs’ YAP signalling respond directly to intrafollicular pressure. It is worth nothing that in *Drosophila*, collagenase has been reported to lower collagen IV contents in the BM^67^, thereby lowering the BM stiffness and pressure in *Drosophila* ovarian follicles^68^. In our study, we observed a similar reduction in tissue pressure with collagenase, potentially due to the combined effect of reduced BM stiffness and decreased TC contractility.

The use of dextran to compress tissues has been widely used in spheroids and organ development^48,69^. With this approach, we successfully determined the effective bulk modulus of follicles to be ∼25 kPa, which translates to an apparent shear modulus of less than 10 kPa, assuming the follicle’s Poisson ratio ranges between 0.2 and 0.45, values that are typically found in tissues. This is consistent with the observation that follicles *in situ* are often found to be deformed by the neighbouring follicles or stroma where the ECM stiffness was reported to be in the range of kPa^18^. Importantly, we found that global compression of follicles leads to cytoplasmic YAP localization and reduced proliferation of GCs, similar to what has been reported for cancer cell spheroids^70^. Here, we propose that the increase in tissue packing may promote GC interactions to activate contact inhibition signals of proliferation^71^.

The striking decrease in TZP number density upon perturbation of TC contractility and BM integrity suggests that a release of tissue pressure directly disrupts the oocyte-GC communications that is essential for oocyte maturation^51^. The negligible impact of actomyosin perturbation or BM disassembly on oocyte size (Figure S5D) indicates that the oocytes in their native state do not experience significant compression. In contrast, global compression by dextran does incur a transient decrease in oocyte size, suggesting that the oocytes are compressible, potentially through dynamic fluid exchange with the surrounding oocyte GCs through gap junctions such as Connexin 37^72,73^.

In this work, we introduce a new approach to quantify follicle growth based on growth rate analysis (Figure 7C). This enables us to uncover an initial size-dependent growth rate followed by a terminal growth rate at maturation phase once the follicles grow past a critical size of 180 µm. This is in marked contrast to cancer spheroids which exhibit a constant growth rate independent of its size (logistic growth)^48^. We found a strong impact of reduced TC contractility on follicle growth at maturation phase which is not due to a difference in the number of cells undergoing apoptosis, mitosis, or telophase. One possibility is that reduced TC contractility and tissue pressure may alter the interphase cell cycle length, tissue packing or cell division pattern that affect the follicle growth rates. Future live imaging studies on intrafollicular dynamics will help to shed light on the interplay between tissue pressure and growth, and reveal the origin of non-exponential, size-dependent follicle growth during early secondary follicle development.

Over the years, there has been growing evidence that 3D compressive stresses modulate tissue dynamics and fate specification in mammalian organ development^69,74^, tumour cell progression^46,75^ and spheroid growth^48^. Our data echo these findings and provide concrete evidence that compressive stress can regulate intrafollicular signalling and follicle growth in early female reproductive processes. Based on our preliminary observation that TCs from aged ovaries generally express little pMLC in follicles compared to those from the young ones (Figure 7H), we speculate that mis-regulated TC mechanics and intrafollicular pressure might contribute to age-associated decline in oocyte quality and anovulation during infertility^76^. Our work therefore provides a new conceptual framework in understanding reproductive biology and ageing, with potential clinical implications in assisted reproductive technology.

### Limitations of the study

One limitation of our study is that the global pharmacological perturbations may incur non-specific effects on the GCs and oocytes, although we have shown that the GCs and oocyte cortex express minimal (or a change of) amount of pMLC expression, respectively (Figure S1D-E). Future work using targeted genetic perturbation may further elucidate the specific functions of TC mechanics during ovarian follicle development. Our study does not determine the exact molecular mechanisms underlying the changes in TZPs and follicle growth upon mechanical stress perturbation. We propose that gap junction dynamics in GCs and oocyte-GC interface might be involved in mechanical signalling. A complete understanding of follicle mechanics and oocyte mechanotransduction would require mapping out the intra-follicle mechanical stress distribution, which may benefit from the potential use of 3D force sensors^79^ and stress inference^80,81^.

## Acknowledgements

We thank Apoorva Shivankar for preliminary experiments on tissue staining in aged mouse ovaries. We are grateful to Brenda Nai Mui Hoon and Chwee Teck Lim for training and usage of atomic force microscopy; Teng Xiang and Yusuke Toyama for training and usage of laser ablation experiments; Xianbin Yong and Cheng Kuang Huang for assistance in traction force microscopy, and the lab of Krystyn Van Vliet for providing the gelatin beads. We thank Jacques Prost for discussions on interpreting AFM data. The Chan lab is supported by the Ministry of Education under the Research Centres of Excellence programme through the Mechanobiology Institute and the Department of Biological Sciences at the National University of Singapore, the Ministry of Education Tier2 grant (T2EP30222-0026) and the Bia-Echo Asia Centre for Female Reproductive Longevity and Equality (ACRLE) at the National University of Singapore. C.J.C. acknowledges the support of the Singaporean Teaching and Academic Research Talent Inauguration Grant (START). We thank Raymond Rodgers, Tetsuya Hiraiwa, and Yuchen Long for providing valuable feedback on our manuscript.

## Author contributions

Project Conceptualization and Design: A.B., C.J.C.; Experiments: A.B., B.H.N., Z.W. S.D., T.B.L., K.T., C.J.C.; Data Analysis, Quantification and Statistical Analysis: A.B., Y.L., C.J.C., Writing: A.B., C.J.C.; Data Interpretation: A.B., Y.L., I.B., C.J.C.; Supervision: C.J.C.

## EXPERIMENTAL MODEL AND STUDY PARTICIPANT DETAILS

### Animals

Mice were group housed in individually ventilated cages with access to water and food under a 12-hr light/12-hr dark cycle. Mouse rooms were maintained at 18-25 °C and 30-70% relative humidity. C57BL/6NTac female mice, aged P25 – P28, were euthanized by carbon dioxide asphyxiation followed by cervical dislocation. ICR female mice, aged 9 weeks and 12 months, was used for preliminary experiments reported in Figure 7F. Ovaries were then dissected from the mice and transferred to an isolation buffer consisting of Leibovitz’s L15 medium (Thermo Fisher) supplemented with 3 mg/ml Bovine Serum Albumin (BSA, Sigma). All mice care and use were approved by the Institutional Animal Care and Use Committee (IACUC) at the National University of Singapore.

## METHOD DETAILS

### Pharmacological treatments

Blebbistatin (Selleck) was used at 5 µM or 20 µM and Y-27632 (Merck) was used at 20 µM to inhibit cell contractility. LPA (Sigma) was used at 20 µM to enhance cell contractility. Large dextran molecules (2 MDa, Sigma) were dissolved in growth medium in varying concentrations to generate varying osmotic pressures^82^. 0.2 mg/mL collagenase was used to disrupt the BM. RGDS (Abcam) was used at 40 µM for to inhibit TC-FN adhesion. Transient perturbations were done for 30 mins; perturbations to check changes in pMLC/FN expression was done for 2-4 hours. Verteporfin (Sigma) was used at 5 µM for 6 hours to inhibit YAP signalling.

### 3D follicle culture

Follicles were mechanically isolated from dissected ovaries under the stereomicroscope attached to thermal plate using tweezers in IB at 37 °C. Growth medium, consisting of MEM-α GlutaMAX (Thermo Fisher) supplemented with 10% Fetal Bovine Serum (FBS, Thermo Fisher), 1% Penicillin-Streptomycin (Thermo Fisher), 1xInsulin-Transferrin-Selenium (Thermo Fisher), and 50 mIU/ml follicle stimulating hormone (Sigma) was prepared. Individual follicles were transferred to each well in a 96-well non-treated plate and cultured in growth medium at 37 °C, 5% CO_2_, 95% humidity overnight. 1% alginate (Sigma) was prepared in phosphate buffer saline (PBS, Gibco) and mixed with growth medium in a 1:1 ratio. Follicles were mouth-pipetted to the 0.5% alginate solution, and hydrogels were formed by pipetting each follicle-alginate mix into the crosslinking medium for ∼2 mins. The crosslinker consisted of 50 mM calcium chloride (Sigma) and 140 mM sodium chloride (1^st^ BASE). Once encapsulated, each gel was placed in 100 µL growth medium inside individual wells of Ultra-Low Attachment 96-well plate (Corning). For long term cultures, half the volume of the growth medium was changed in each well every two days. The follicles were removed from the alginate hydrogels after four days using 10 IU/mL alginate lyase (Sigma) at 37 °C for 10-15 mins.

### Bead-follicle assays

Follicles were placed in the follicle medium filled with red or green cytoplasmic membrane dye (Cellbrite) for 1-2 hours to label the outer theca cells. They were then washed thrice before being transferred to 200-μl droplets of follicle medium filled with the gelatin beads (kind gifts from Krystyn van Vliet’s lab) in a 35 mm glass-bottom dish (Cellvis) covered with mineral oil (Sigma). The beads and follicles are manipulated to position them in contact and cultured in an incubator with a humidified atmosphere supplemented with 5% CO_2_ at 37 °C for up to 2 hours. Time-lapse imaging for bead-follicle fusion was performed on a Zeiss LSM 710 confocal microscope with an onstage incubator using 40×/NA 1.2 W Corr objective and Zen 2012 LSM software with 488 nm and 633 nm lasers. Image stacks were acquired at 90 mins intervals with 4 μm z-steps. For bead-follicle experiments performed in the presence of blebbistatin, the follicle medium (volume increased to 400 μl) was not covered with mineral oil.

### Tissue sectioning

Ovaries were fixed in 4% paraformaldehyde (PFA, Santa Cruz Biotechnology) at room temperature (RT) for an hour. The fixed ovaries were washed in washing buffer (WB, 1% BSA in 1X PBS) thrice before being embedded into 4% low-melting point agarose (Thermo Fisher). The embedded tissue was sliced into 100 µm thick tissue sections using a vibratome (Leica) in PBS at 0.05 mm/s speed and 1 mm amplitude.

### Immunofluorescence staining

Isolated follicles were fixed in 4% PFA at RT for 30 mins and washed with WB thrice before immunostaining. Fixed samples were incubated in blocking-permeabilizing solution (3% BSA and 0.03% Triton X-100) at RT for 2-4 hrs, followed by incubation at 4 °C in primary antibodies diluted in the blocking solution overnight. The tissues were washed 5 times in WB and incubated in secondary antibodies diluted in the washing buffer for 4 hrs at RT. They were washed thrice in WB before mounting. The ovarian slices were mounted into ProLong Gold antifade mountant (Thermo Fisher) and left to cure overnight at RT, whereas isolated follicles were mounted into SlowFade Gold antifade mountant (Thermo Fisher) prior to imaging.

Primary antibodies used were rabbit anti-phospho myosin light chain 2 (Ser19) (Cell Signaling Technology, 1:100), rabbit anti-fibronectin (Abcam, 1:100), rabbit anti-Ki67 (Cell Signaling Technology, 1:100), rabbit anti-phospho histone H3 (Cell Signaling Technology, 1:100), rabbit anti-cleaved caspase 3 (Abcam, 1:100), and mouse anti-YAP (Abnova, 1:100). Alexa Fluor

488 labelled anti-mouse (Invitrogen, 1:500) and Alexa Fluor 546 labelled anti-rabbit (Invitrogen, 1:500) was used as secondary antibodies. DNA was stained with DAPI (Sigma, 2 μg/mL) and F-actin was stained with either Alexa Flour 488-labelled phalloidin (Invitrogen, 1:1000) or Alexa Flour 633-labelled phalloidin (Invitrogen, 1:300).

All fixed samples were imaged with Nikon A1Rsi confocal microscope with NIS Elements. Isolated follicles were imaged using Apo 40×/1.25 WI λS DIC N2 objective at 4 µm z-slices. Tissue slices were imaged with Plan Apo VC 20×/0.75 DIC N2 and stitched with 10% overlap using lasers 405 nm, 488 nm, 561 nm, 640 nm.

### EdU incorporation assay

Follicles were incubated with 50 µM EdU (EdU Staining Proliferation Kit, abcam) for 2 hours under optimal growth conditions. They were washed with WB and grown under different conditions for 30 mins. The samples were fixed and permeabilized. The EdU reaction solution was prepared as per the manufacturer’s specifications. The follicles were incubated in the reaction solution for 2 hours at RT in the dark. They were washed and incubated with DAPI for 2 hours at RT in the dark before being washed and then mounted for imaging.

### Atomic force microscopy

#### Sample preparation and setup for single cell and follicle indentation

Wafer for the PDMS microwells was designed by the lab. PDMS and cross linker were mixed in 10:1 ratio and degassed before transferring to the wafer. The PDMS mixture was degassed again and cured at 80℃ for two hours. The PDMS mould was removed and trimmed into working size. To fabricate the microwells, PDMS mixture was transferred onto the glass bottom dish (WPI FD35) and the trimmed PDMS mould was placed inverted on top and cured at 80℃ for two hours. The mould was then removed and the PDMS microwells were used for AFM. The microwells were filled with growth medium and left in the 37 °C incubator for at least 30 mins. Freshly isolated follicles were transferred to medium and left to stabilize under optimal growth conditions before being indented.

The NanoWizard 4 BioScience (JPK Instruments AG) mounted on an inverted microscope (Olympus IX81) with a 10x objective was used. Polydimethylsiloxane (PDMS) microwells with 100 µm, 130 µm, 150 µm diameters at 50 µm spacing and 80 µm depth were fabricated to house the follicles during AFM experiments.

A pyramidal tip on Bruker MLCT-D cantilever (0.03 N/m spring constant) was used to measure effective TC surface tension. A polystyrene particle (45 μm) on silicon nitride cantilever (Novascan Technologies, 0.35 N/m spring constant) was used to measure bulk tissue mechanics. Measurements were conducted with a constant speed of 5 μm/s, with a loading force of 10 nN (tip) or 15 nN (bead) in a 10 μm by 10 μm area. Both sensitivity and spring constant were calibrated using contact-based approach prior to each experiment. Follicle diameters and effective tip radius were determined from the brightfield images.

### Primary cell isolation

Primary ovarian cells were isolated based on protocols adapted from Tingen et al.^38^. In brief, freshly isolated ovaries were poked in IB by a needle under the stereomicroscope to release the GCs till intact follicles were no longer observed. This solution was centrifuged at 94g for 5 mins. The pellet was washed twice and resuspended in McCoy’s 5A medium (Gibco) supplemented with 5% FBS and 1% Penicillin-Streptomycin to yield primary GCs.

A digestion buffer comprising of 0.05 mg/mL activated DNAse I (DNase I with HBSS in a 1:1 ratio, Merck), 10 mg/mL Collagenase (Thermo Fisher) and 40% Medium 199 (Gibco) was freshly made. The remaining tissue fragment after mechanical disruption containing theca cells (and stromal cells) was washed and transferred to the digestion buffer (200 µL per ovary). This was incubated at 37 °C for 1 hr mixing gently every 15 mins using a pipette. Once completely dissolved, the solution was centrifuged at 94g for 5 mins. The pellet was washed and resuspended in supplemented McCoy’s medium. Cells were then seeded onto 6-well plates in the growth medium and incubated at 37 °C, 5% CO_2_ and 95% humidity for at least a day before further experiments.

### Traction force microscopy (TFM)

#### a) Preparation of TFM substrates

Glass coverslips were cleaned with 2% Hellmanex III, washed with water, and blow dried with nitrogen before silanization. They were incubated in the silanization solution, 2% trimethoxysilyl propyl methacrylate (TMPMA, Sigma) and 1% glacial acetic acid in absolute ethanol for 10 mins, rinsed with ethanol, blow dried with nitrogen, and incubated at 120 °C for an hour.

An aliquot of 3 µl of 100 nm fluorescent microspheres (F8810, Invitrogen) was added to 10 ml of milliQ water and sonicated for 10 mins. The bead solution was filtered by a 0.22 µm syringe filter (Sartorius) into 500 µL of 500 mM MES buffer (pH 6.0). A master polyacrylamide solution was made by mixing 200 µL of 40% polyacrylamide solution (Biorad), 200 µL of 2% bis-acrylamide solution (Biorad), 1.5 µL TEMED (Sigma), and 582.5 µL of the bead solution. 16 µL of 10% ammonium persulphate (Biorad) was added to the master mix, and 100 µL droplets were dispensed on a clean parafilm strip. The silanized coverslips were gently placed on the gel droplets such that the gel covered the whole glass area, and the polyacrylamide was allowed to gel for 30 mins. The coverslips with the gel were removed from the parafilm by floating water at the bottom of the gel and placed gel side facing upwards in PBS at 4 °C overnight after washing with 1X PBS thrice.

The gel was soaked in 0.1 M HEPES (1^st^ BASE, pH 7.4) for 30 mins. A 0.02 mg/mL solution of sulfosuccinimidyl 6-(4’-azido-2’-nitrophenylamino) hexanoate (sulfo-SANPAH, 803332, Sigma) was prepared in anhydrous DMSO. 1 µL of this sulfo-SANPAH solution was added to 20 µL of 0.1 M HEPES (pH 7.4). This solution was added to the polyacrylamide gel surface after discarding the HEPES that was soaking the gel. Mechanical agitation by a silicone block was done to ensure that sulfo-SANPAH was distributed uniformly. The gels were UV treated in the UV-KB9 (KLOE, France) at 8% power for 5 mins, and washed twice in 0.1 M HEPES (pH 7.4). The process was repeated with a fresh solution of sulfo-SANPAH. The gels were washed twice with 0.1 M HEPES (pH 7.4) and once with 1X PBS.

A 100 µg/mL solution of collagen I (Sigma) was prepared in 1 X PBS. 250 µL of this solution was added to the gel and incubated in the dark for 2 hours at room temperature with intermittent mixing with a pipette to avoid clumping. The gels were washed thrice with 1 X PBS and stored in PBS before seeding of cells.

#### b) TFM setup

A spinning disk-confocal microscope with a Yokogawa CSU-W1 scanner unit (Yokogawa Electric, Japan), an iLAS laser launcher (Gataca Systems, France), and a sCMOS Camera (Prime 95B 22 mm, Teledyne Photometrics, USA) attached to a Nikon Ti2-E was used. Images were acquired by a 40x water immersion objective (CFI Apo LWD 40XWI λS N.A. 1.15, Nikon) at z-steps of 0.275 µm with the help of MetaMorph advanced acquisition software (Molecular Devices, USA). Coverslips with the polyacrylamide gels were placed in stainless steel cell culture vessels and 500 µL of supplemented McCoy’s cell culture medium was added. Custom-made lids were used to control temperature, CO_2_, and humidity while imaging. A z-stack of the fluorescent labelled beads was captured when the cells were adhered to the gel. 1% sodium dodecyl sulfate was prepared in the supplemented culture medium and 100 μL of this was added to the gel. A second z-stack of the beads was acquired with the same settings after 30 mins.

### Scanning electron microscopy (SEM)

Ovaries were fixed with 2% PFA and 3% glutaraldehyde (GA) overnight at 4 °C. They were washed thrice with PBS for 5 mins each. Samples were incubated in 1% osmium tetraoxide (OsO4) with 1.5% potassium ferrocyanide in PBS for 1 hour on ice and then washed thrice with distilled water for 5 mins each. The samples were then placed into 1% thiocarbohydrazide (TCh) in distilled water for 20 mins at room temperature and washed thrice with distilled water for 5 mins each. The samples were then placed into 1% OsO4 in distilled water for 30 mins at room temperature and washed thrice with distilled water for 5 mins each. They were next incubated with 1% uranyl acetate (UA) in distilled water overnight at 4 °C and washed thrice with distilled water for 5 mins each. 0.02 M lead nitrate and 0.03 M aspartic acid were mixed, and pH was adjusted to 5.5. The samples were kept in lead aspartate solution for 30 min at 60 °C in the oven, and again washed thrice with distilled water for 5 mins each. Tissues were dehydrated with ethanol, increasing gradually from 25%, 50%, 75%, 95% and 100%, with 10 mins in each solution on ice before washing with acetone twice for 10 mins each on ice. For resin infiltration, the samples were placed in 1:1 acetone-araldite resin mixture for 30 mins and then 1:6 mixture overnight. They were then transferred to pure araldite for 1 hour in a 45 °C oven. This was done thrice before they were transferred into embedding mould with pure araldite and cured for 24 hours in a 60 °C oven.

The embedded samples were then sectioned using Diatome diamond knife with the Leica UC6 ultramicrotome and 100 nm ultrathin sections were collected onto silicon wafers. SEM Imaging was done with Thermofisher FEI Quanta 650 FEG-SEM where large area montage scans were acquired with MAPS 2.1 software using the backscatter mode (vCD detector) at 5 kV, 5 mm working distance (WD).

### Laser ablation

Follicles were stained with EGFP-CNA35 at 8 µM for 2 hours in follicle growth medium to label the BM. Laser ablation experiment was performed on a NikonA1R Multiphoton laser scanning confocal microscope with an Apo 40x/ NA 1.25 WI λ S DIC N2 objective lens. UV laser with 355 nm, 300 ps pulse duration and 1 kHz repetition rate (PowerChip PNV-0150-100, team photonics) was irradiated to the BM in the follicle equatorial plane for 5 secs at 300 nW laser power at the back aperture. For YAP experiments, the follicles were fixed within 2-5 mins after ablation. For velocity calculation, transmitted light (TD) and EGFP channel images were obtained every 2 secs for 10 mins.

## QUANTIFICATION AND STATISTICAL ANALYSIS

### Quantification of basal TC-BM matrix thickness

Line-scans (∼1.5 µm) were drawn perpendicular to the matrix between the basal TCs and BM in the SEM images, spaced 20 µm away from each other. Plot profiles were plotted; and the x-coordinates of the start and end of the “bright” matrix was noted from each intensity profile. The width of each profile was calculated by subtracting the x-values and averaged over all widths obtained from all line-scans in a follicle.

### Quantification of pMLC and FN expression

Using FIJI, the z-plane where the oocyte diameter was the largest in the entire image stack was selected. A polygonal selection tool on the phalloidin stained actin channel was used to mark the TC layer and the GC layers in follicles. The selection was overlaid on the pMLC/FN channel, and the mean intensity of the selection was measured. A 60 x 60 pixels area was demarcated in the same z-plane in the background of the image using the rectangle selection tool and the mean intensity of this area was measured. The ratio of the mean intensity of the signal to that of the background was termed as pMLC or FN expression. The ratio of the TC to GC mean intensity was termed as ratio (pMLC or FN intensity).

A segmented line tool was used to mark the oocyte cortex. The selection was overlaid on the pMLC channel, and the mean intensity was measured. By dividing this value by the mean intensity of the background, the ratio (pMLC intensity) at the oocyte cortex was computed.

### Quantification of AFM-based indentation

The details of AFM-based analyses are explained in Supplementary Information.

#### a) Analysis of effective TC surface tension and effective follicle pressure

There is a linear regime of force-displacement relationship for indentation depth within 100-700 nm. The linear coefficient, here, is related to the hydrostatic pressure exerting at this shell^83,84^ (details in Supplementary Material Sec. 1.a). Assuming a material homogeneity at the scale of follicle size, this pressure could be regarded as an effective hydrostatic pressure of follicle, and its surface tension (mostly contributed by theca cells) was then inferred from Laplace law as *σ*_∞_ = *PR_f_*/2, where *R_f_* is the follicle radius.

#### b) Analysis of tissue elasticity and hysteresis

For indentation depth within 1∼5*μ*m, which is much smaller than the of probing bead and follicle radius, we used a Maxwellian viscoelasticity model to extract the pure elastic parts of approach and retraction curves. The pure elastic force-indentation curve was then fitted by simple Hertz model for a bead tip. Hysteresis was calculated as the area under the curve between approach and retraction plots.

### Quantification of local parallel velocity

PIV analyses were implemented onto the time-lapse images obtained after laser ablation using openPIV in Python. The range of local area around the ablation point was determined by the GC layer thickness. The velocity of GC cells flowing away from the follicle centre was quantified as the mean velocity in this local range projected along the direction pointing from ablation point to the follicle centre.

### Quantification of bulk moduli for follicles and beads

A polygonal selection was drawn on the edge of the follicle/bead using FIJI. The Fit Ellipse option was used to measure the major (a) and minor (b) axes of the selection. Volume of the

selection was calculated by 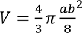. The difference (Δ*V*) between the initial and final volume was computed for every osmotic pressure and the ratio of the difference to the initial volume at each osmotic pressure was plotted with the corresponding osmotic pressure 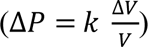, where *k* is the bulk modulus. Curves with negative data points were removed and the average curve was generated. The linear part of the average plot, between 0 to 5 kPa, was fitted to a straight line with the intercept fixed at 0. The slope was measured; the inverse of the slope was calculated and termed as bulk modulus.

### Quantification of TC compressive stress

The z-plane of the image stack (captured at t = 0 hr) with the maximum bead diameter was determined. A maximum intensity projection was obtained from 5 slices (the max. bead diameter z-plane, 2 slices before and 2 slices after that). The outline of the bead was demarcated, and the volume of the bead is measured. The same was done for the image stack at t = 40 hours, and the change in volume between these two timepoints was calculated. The compressive stress was then computed by multiplying the relative volumetric change of the beads to its bulk modulus.

### Quantification of traction stress

Images of the same cells with and without beads (after SDS washing) were stacked to create an image pair using FIJI. An ImageJ plugin, Align Slice, was used to align for any drift away from the cell boundary. The cell boundary was noted from the corresponding brightfield image. The bead displacement field and magnitude were calculated using the PIV plugin using the same iteration scheme (128/256 for 1^st^ pass, 64/128 for 2^nd^ pass, and 32/64 for 3^rd^ pass). The threshold, or the cross-correlation coefficient was set at 0.60. The traction stress field and magnitude were computed by the FTTC plugin using 32 kPa as the stiffness, 0.5 as the Poisson ratio, and 9×10^−11^ as the regularization factor. The stress fields were read out as images in FIJI; the cell brightfield images were used to outline the boundary of the cells and stress clusters within each cell were identified by overlaying the cell ROI using Particle Analysis plugin. The cluster ROIs were then overlaid on the stress magnitude images; average and maximum stress from each cluster was then measured.

### Quantification of follicle volume upon transient perturbations

Freshly isolated follicles (one in each well) were placed in growth medium, and images of each follicle were immediately captured. They were incubated at 37 °C for 30 mins and images of each were acquired again. They were then transferred to medium containing Blebb/Y-27632 and images were captured. The follicles were imaged again after 30 mins and transferred to normal growth medium. Images were taken instantly and then after incubating for 30 mins at 37 °C. Each follicle, thus, could be tracked over six images. Follicle volume was calculated as mentioned in the previous section. The ratio between follicle volume at t = 30 hr and t = 0 hr for each condition (control, treatment, recovery) was termed as ratio (Follicle volume).

### Quantification of GC proliferation and YAP signalling

The z-plane of the image stack (isolated follicles and tissue slices) with the maximum oocyte diameter was determined in FIJI. The number of DAPI-stained nuclei and Ki67-labelled nuclei in this layer was counted separately. The ratio between the Ki67+/DAPI+ was calculated, and the value was termed Ki67 positive. The same approach was taken for the EdU proliferation analysis.

Nuclei and cytoplasm of each GC were identified using the DAPI and DAPI/Phalloidin overlay respectively. A 2×2 pixels selection was drawn each on the nucleus and its corresponding cytoplasm. These selections were overlaid on the YAP channel and the mean intensity was measured for each selection. The ratio of the nuclear to cytoplasmic selections for each cell was calculated and plotted.

### Quantification of number of Transzonal projections (TZP)

The z-plane of the image stack with the maximum oocyte diameter was determined. The intensity of the background of the sample was measured at this plane. A segmented line was drawn on the zona pellucida surrounding the oocyte using the Line ROI tool in FIJI. The length of this line was measured. The intensity of the actin-labelled image was plotted as a function of the length of the line using Plot Profile in FIJI. The data was used to count the number of peaks above the background value using Origin2021b. The ratio of the number of peaks to the length of the line was termed as number density.

### Quantification of interstitial gap within follicles

A pixel was detected as interstitial if its intensity was below the background noise level in all three channels of DAPI, Actin and YAP. A cluster of connected interstitial pixels was recognized as one interstitial gap. Each cluster has an area of *A* and perimeter length of *P*, the shape of each cluster is quantified as the ratio, which is 1 for a sphere and larger than one for an elongated shape. All algorithms were developed with OpenCV in python.

### Quantification of GC number

Cell nuclei segmentation was performed on the DAPI channel using a Birch clustering algorithm. Pixels close to one cluster seed was identified as one cell. Image processing codes were implemented by Python. Birch algorithm was implemented through the scikit-learn module. A heurist parameter-tuning method was applied to the algorithm without assigning a cluster number to search for the optimal cluster (cell) size that would minimizes the within-cluster-sum (wss) score. Then, the algorithm was looped over a reasonable range of cluster no. using this optimal cluster size to find the optimal cluster (cell) number that had the lowest wss score. The number of clusters in CC3 and PHH3 channels (number of cells with positive signals) are found by the same clustering algorithm with the optimal cluster size obtained in the DAPI channel.

### Quantification of follicle growth

Follicle diameters *D* were measured by length measurements tools in FIJI. Growth rate *γ* of follicles was defined as the change of diameter d*D* over a period of time d*t*, normalized by the diameter: 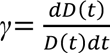, which is equivalent to *d* ln *D*(*t*)/*dt* when d*t* is small. In practice, we calculated the growth rate of a follicle with diameter *D* at time *t* from the discrete time evolution as (ln(*D*(*t*)) − ln (*D*(*t* − Δ*T*)) /Δ*T*, where Δ*T* is the time interval between two consecutive time points and Δ*T* is usually 1 day or 2 days. The growth rate for diameters >180 µm was used to plot G(r) at the maturation phase.

### Statistical analysis

All graphs and statistical tests were created using Origin 2021b. N represents the number of independent experiments and n represents the total number of follicles/tissues in the representative data shown in figures. The data was tested for significance using Mann-Whitney U test when ns: p>0.05, *: p<0.05, **: p<0.01, and ***: p<0.001.

## Supplementary Information

### Supplementary Methods

AFM-based indentation analyses, a schematic illustration shown in Fig. IA.

**Figure I:**
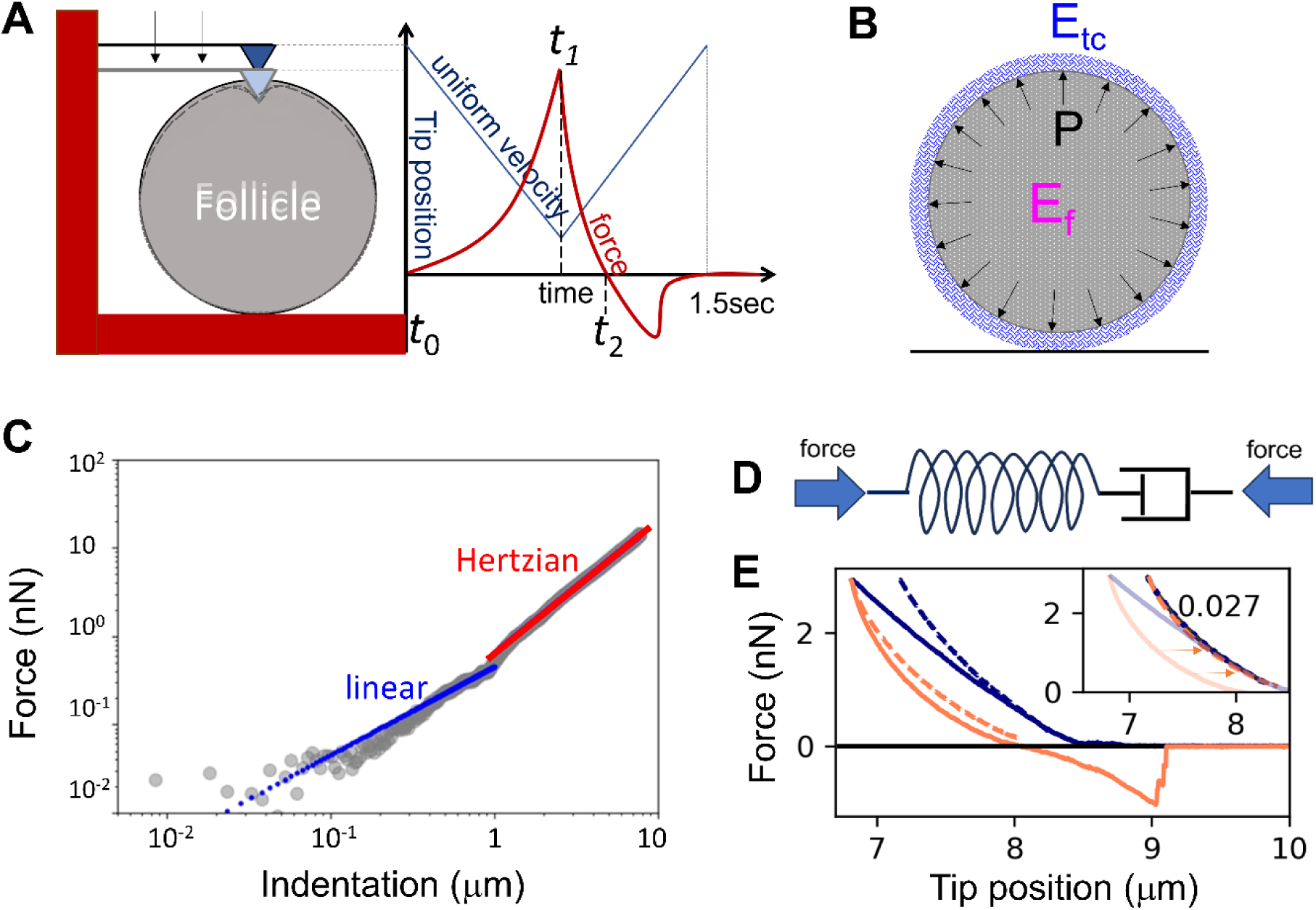
Atomic Force Microscopy (AFM) and the time evolution of the tip position and measured force from the cantilever. (A) The cantilever (example of a pyramidal tip is shown here) is moving with uniform velocity downward until the force resisting the tip of cantilever reaches the pre-set maximum force value or the tip position reaches the pre-set maximum range. The total time duration is about 1.5 second. (B) Model of follicle. (C) A typical two-regime force-indentation relationship with a bead tip of radius 22.5 μm, where a linear regime at hundreds nm scale and a non-linear regime at micron scale following Hertzian contact model were found. (D) Maxwellian model of viscoelasticity. The elastic component (left) and a viscous dashpot (right) are connected in series, along which the elastic force and the viscous force is balanced, while the total strain, which is characterized by the tip displacement, is the summation of the contraction two components. (E) A typical approach (straight navy) and retraction (straight orange) force-tip position curve generated from one AFM measurement. Dashed curves are the inferred elastic part of the curves based on a Maxwellian model. Inset: the two inferred elastic curves exhibit similar force-position profiles with a mean squared percentage difference 2.7%, which is drastically reduced from the original difference ∼28%.

#### a) Analysis of follicle surface tension and pressure

Let’s assume the follicle is a pressurized elastic ball covered by an elastic shell of theca cells (Fig. IA-B) with a thickness *h*, elastic modulus *E*_tc_, a homogeneous radius of curvature *R_f_* and standard Poisson ratio *ν_tc_*. Then deformation caused by a poke from a pyramid tip is composed mainly of the deformation at the tip side *δ*_tip_ and the deformation at the bottom side *δ*_bot_. The force balance at each side reads *F*_*ela*_(*δ*) + *F*_*p*_(*δ*) = *F*(*δ*), where *F*_*ela*_is the elastic force (including bulk compression and bending) that resists the external poke *F*(*δ*), and *F*_*p*_(*δ*) is the force due to the hydrostatic pressure. As the elastic shell of theca cells is much thinner than its radius of curvature, a deformation *δ* smaller than the thickness of *h* renders the force linear to the deformation (Fig. IC), which could be explained by the theory for a compressed thin elastic shell^1,2^. If the radius of contact area *s_c_* is much smaller than the characteristic length scale of bending *l_b_∼*(*R_f_ h*)^0.5^, the force *F* = *K*_1_*δ*, where *K*_1_ is the shell stiffness mainly contributed by shell bending; otherwise, if the radius of contact area *s_c_* is comparable or larger than the bending length scale *l_b_*, we estimate *F* ∼ *K*_2_*δ*, where *K*_2_ is the shell stiffness contributed by geometric stretching of shell due to pressure with bending neglected.

In Fig. 1E, we probed the follicle depth up to 2∼2.5 μm with a pyramid tip of 20 nm. At the top side, and the contact radius of the nanotip is far smaller than the bending length scale *l_b_*, which is on the micron scale, therefore, we have *F* = *K*_1_*δ*_tip_ at the tip side. At the bottom side, the contact radius *s_c_ ∼* (2*δR_f_*)^0.5^, close to *l_b_*, therefore we have *F* ∼ *K*_2_*δ*_bot_ at the bottom. Finally, we estimate the force *F* in relation to total indentation *δ* as *F* = *Kδ*, where the apparent stiffness *K* follows 1/*K*= 1/*K*_1_+1/*K*_2_, for the *δ* in the linear regime.

The force balance equations for the shell in polar coordinates (also see^1,2^) are:

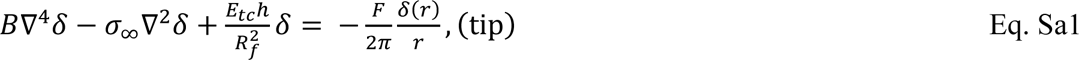

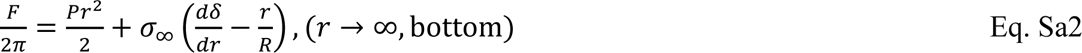

where *r* is the horizontal distance of points on the shell to the central axis, *B* is the bending modulus of shell, *σ*_∞_ = *PR*_*f*_/2 is the natural shell surface stress (Laplace’s Law holds where *r* is far from the central axis). We then obtain the shell stiffness *K*_1_ from the Eq. Sa1 as

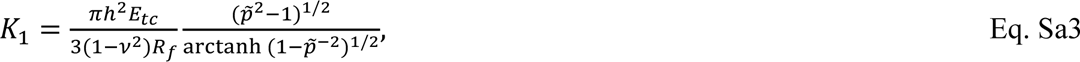

where *p̃* is a dimensionless pressure

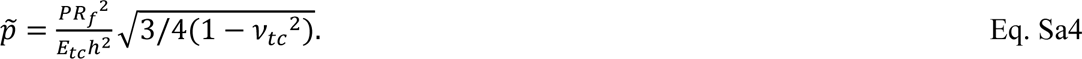

For 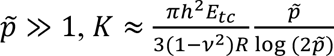, which depends both on the pressure and shell elasticity; for 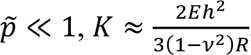, not relative to pressure.

From Eq. Sa2, we obtain *K*_2_ as

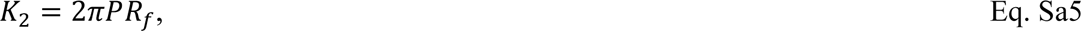

which merely depends on the pressure, regardless of the contact geometry. One can easily find that *K*_1_ is much smaller than *K*_2_, therefore the indentation is dominated by the deformation at the tip side.

The apparent shell stiffness *K* is measured from the linear force-indentation regime *δ* from 100 nm to 700 nm. The elasticity *E_tc_* is the apparent elasticity fitted using Sneddon model (see methods below) for the nonlinear regime *δ>*1 *μ*m from the approach curve. Assuming the elastic shell thickness *h* =2 *μ*m and its Poisson ratio *ν_tc_* ∼0.45, pressure was then obtained at ∼20 Pa, which corresponds to a dimensionless pressure *p̃*∼10 ≫ 1. This indicates that the system is in the pressurized condition and therefore the value of pressure obtained is valid. We then obtained the natural shell tension *σ*_∞_ = *PR*_*f*_/2, which is ∼0.55 mN/m for control, 0.9 mN/m for LPA and 0.4 mN/m for Blebbistatin.

In Fig. 3A-D (main text), we probed follicles by a bead tip of radius 22.5 μm. The contact radius at both the tip and the bottom sides is close to the bending length. Neglecting the bending term, the apparent shell stiffness *K* ∼ *K*_2_/2, from which we understand that deformation from both sides contribute equally to the indentation. From Eq. Sa5, we fitted the shell surface pressure, and they have similar values with the ones measure from a pyramid tip.

#### b) Analysis of tissue viscoelasticity

##### b.1#A Maxwellian viscoelasticity model

For indentation *δ* > 1*μ*m, the force-indentation curves show superlinear powers (Fig. IC). This is because the resisting force is contributed mainly by the viscoelastic bulk deformation of the follicle surface materials (including the theca cells, basement membrane and basal granulosa cells) and the linear contribution due to the hydrostatic pressure is negligible. To extract the viscoelasticity of these parts, we use a Maxwellian model of viscoelastic fluids (Fig. ID). The force *F* at the tip is balanced in the elastic part and viscous part in series; and the displacement *δ* is the summation of the elastic and viscous counterparts. Hence, the force measured by the tip *F* and tip displacement *δ* in approach and retraction processes in the positive force region are assumed to obey the following evolution over time *t*, respectively:

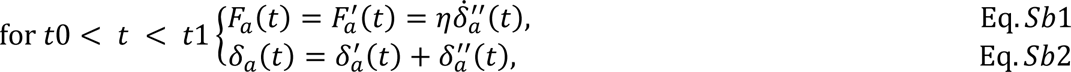

and

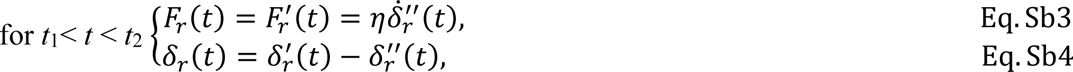

with boundary conditions *F*_*a*_(*t*_0_) = *F*_*r*_(*t*_2_) = 0, *F*_*a*_(*t*_1_) = *F*_*r*_(*t*_1_), *δ*_*a*_(*t*_0_) = 0, *δ*_*a*_(*t*_1_) = *δ*_*r*_(*t*_1_).

The elastic force, denoted by *F*^′^, is related with the elastic displacement *δ*^′^ using a Hertz model as 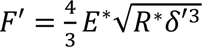 (for bead tip), or a Sneddon model 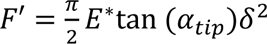 (for pyramid tip), where *E*^∗^and *R*^∗^are the effective elastic modulus and effective radius. Since the bead is relatively rigid, *E*^∗^ ≈ *E*/(1 − *v*^2^), where *E_f_* is the follicle elasticity to be measured and *ν* is set as 0.45 the follicle’s Poisson’s ratio. The effective radius is calculated as 1/*R*^∗^ = 1/*R*_*f*_ + 1/*R*_*b*_, where *R*_*f*_ is the follicle radius and *R*_*b*_the bead tip radius.

Substituting Eq. Sb1(or Sb3) into Eq. Sb2(or Sb4) gives us the elastic component displacement which can be related to force as

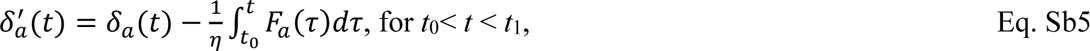

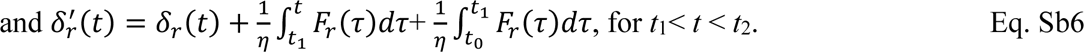

The viscous force, 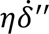, is assumed as linear to the shrinkage rate of the viscous damper 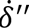, with the viscosity coefficient *η* in the unit of Ns/m to be measured from the data. The displacement of the viscous components is calculated as 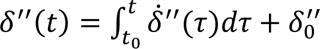, where 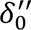 is zero for *t*_0_.

As *F*(*t*_2_) =0, the elastic component at *t*_2_ also has zero displacement. From Eqs. Sb1-2, the distance between the tip position *δ_r_*(*t*_2_)-*δ_a_*(*t*_0_) is derived to be

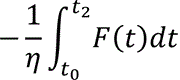

From the data, we could directly measure *δ_r_*(*t*_2_)-*δ_a_*(*t*_0_) and thus obtain the Maxwellian viscosity of the system.

Substituting this viscosity into Eq. Sb5, we calculated the elastic component displacement in relation to *F_a_* and used the corresponding bulk models to fit the elasticity of the follicle *E*.

We can also substitute the viscosity value into Eq. Sb6 to obtain another corresponding elastic component displacement in relation to *F_r_*. A verification on this assumption of Maxwellian viscoelasticity is to compare the two elastic force-displacement curves from Eq. 5 and Eq. 6. Consequently, the two curves were sharing a similar profile (Fig. IE, dashed) with a mean squared difference dropped from the original value of 28.1% to 2.7%, suggesting a good performance of Maxwellian model.

Note that the indentation value in the elastic force-indentation curve is contributed by both the tip and the bottom side. For the pyramid tip measuring mainly the theca cell layer elasticity 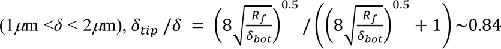, *i.e*., deformation at the tip side contribute dominantly. For the bead tip measuring the thicker layer composed of theca and basal granulosa cells 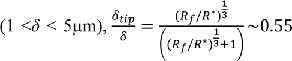, *i.e*., deformation at both sides contribute almost equally. After extracting the tip fraction of indentation, we finally fit the elasticity from the pure elastic force-indentation curve at the tip side for the theca cell layer (pyramid tip) and for the thicker tissue bulk (bead tip), respectively.

##### b.2 Hysteresis

The energy done by the tip to follicle during approach is 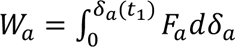, while the energy released by the follicle during retraction 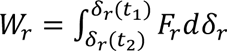. Work loss is defined as the energy dissipated over the whole loop of approach and retraction:

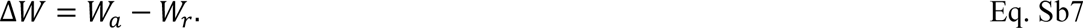

The value of work loss can be directly measured from the approach and retraction curves of force and tip positions.

The hysteresis mentioned in the main text is then calculated as Δ*W*/*W*_*a*_, which is the energy lost in the whole indentation process normalized by the total work done in the approach phase. As the elastic components do not dissipate energy during the processes, the work loss is merely contributed by the viscous component of the system:

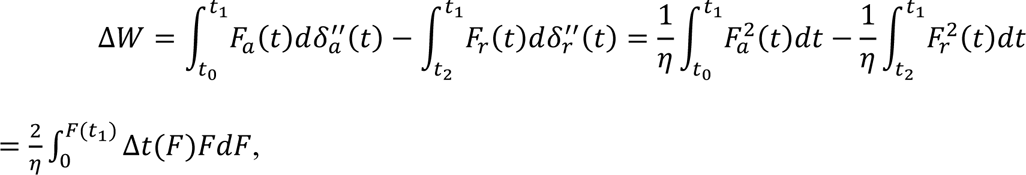

where Δ*t*(*F*) = *t*_2_ − (*t*_*r*_(*F*) + *t*_*a*_(*F*)) is related with the two times at the same value of *F* in approach and retraction processes.

## Supplementary Figures

**Figure S1:**
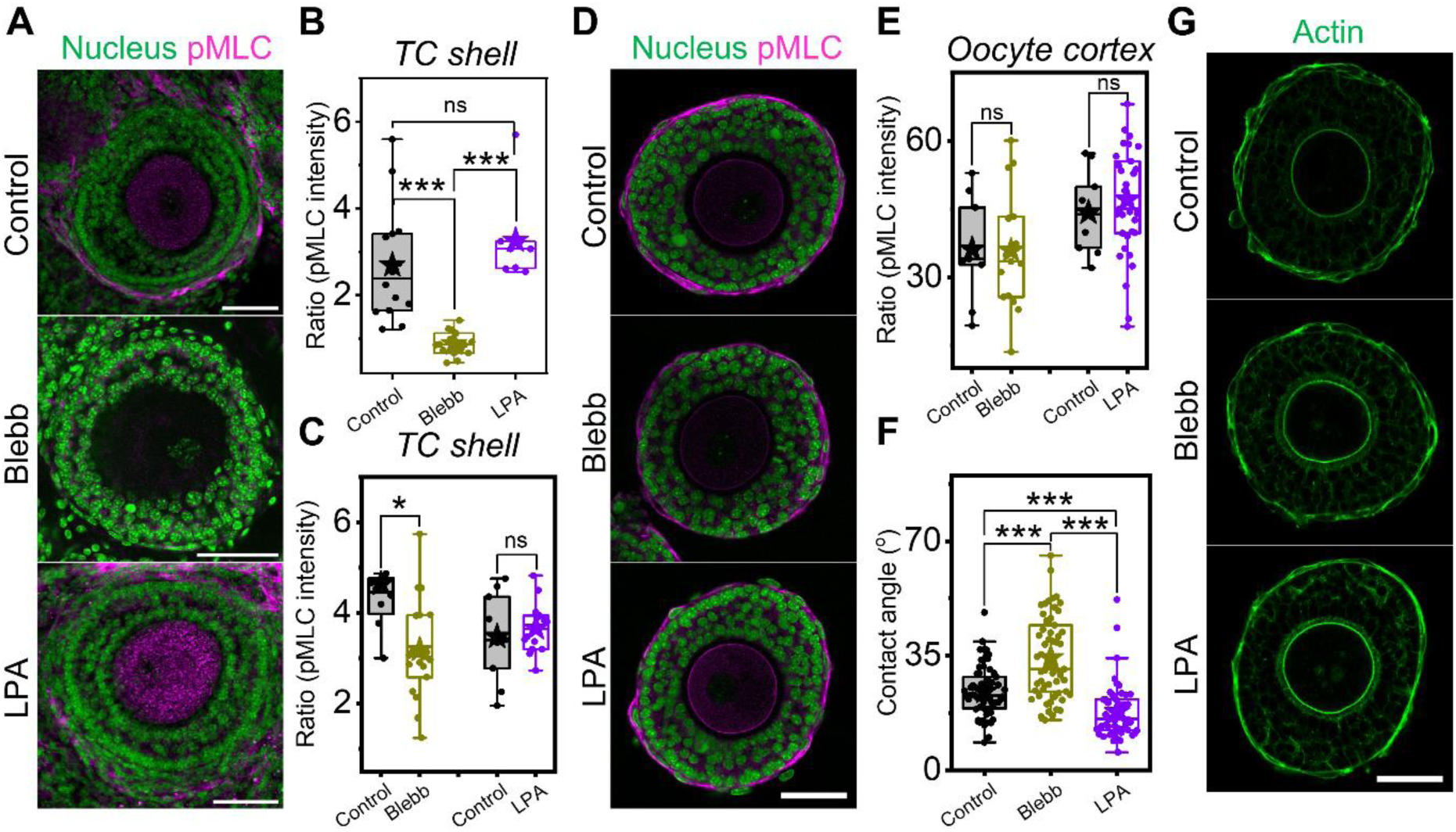
Actomyosin perturbation impacts TC cortical tension but not the oocyte cortical tension, related to Figure 1. A) Representative images of tissue slices labelled with DAPI (nucleus, green) and immuno-stained with pMLC (magenta) in control, Blebb, and LPA-treated samples. Scale bar: 50 µm. B) Boxplots of ratio (pMLC intensity) at TC shells in various conditions in ovarian tissue slices (*in situ*). N = 2, n = 15 follicles. C) Boxplots of ratio (pMLC intensity) at TC shells of isolated secondary follicles in various actomyosin perturbations. N = 2, n = 10-14 follicles each. D) Representative images of isolated follicles (*ex vivo*) in control, Blebb, and LPA-treated samples stained with DAPI (nucleus, green) and immuno-stained with pMLC (magenta). Scale bar: 50 µm. E) Corresponding boxplots of ratio (pMLC intensity) at oocyte cortex in various conditions. N = 2, n = 12-14 follicles each. F) Boxplots of contact angle of TCs on follicles in various conditions. N = 2, n = 12-14 follicles each. G) Representative images of isolated follicles stained with Phalloidin (actin, green) in various conditions. Scale bar: 50 µm. Significance was determined by Mann-Whitney U test. ns: p > 0.05; * p < 0.05; *** p < 0.001.

**Figure S2:**
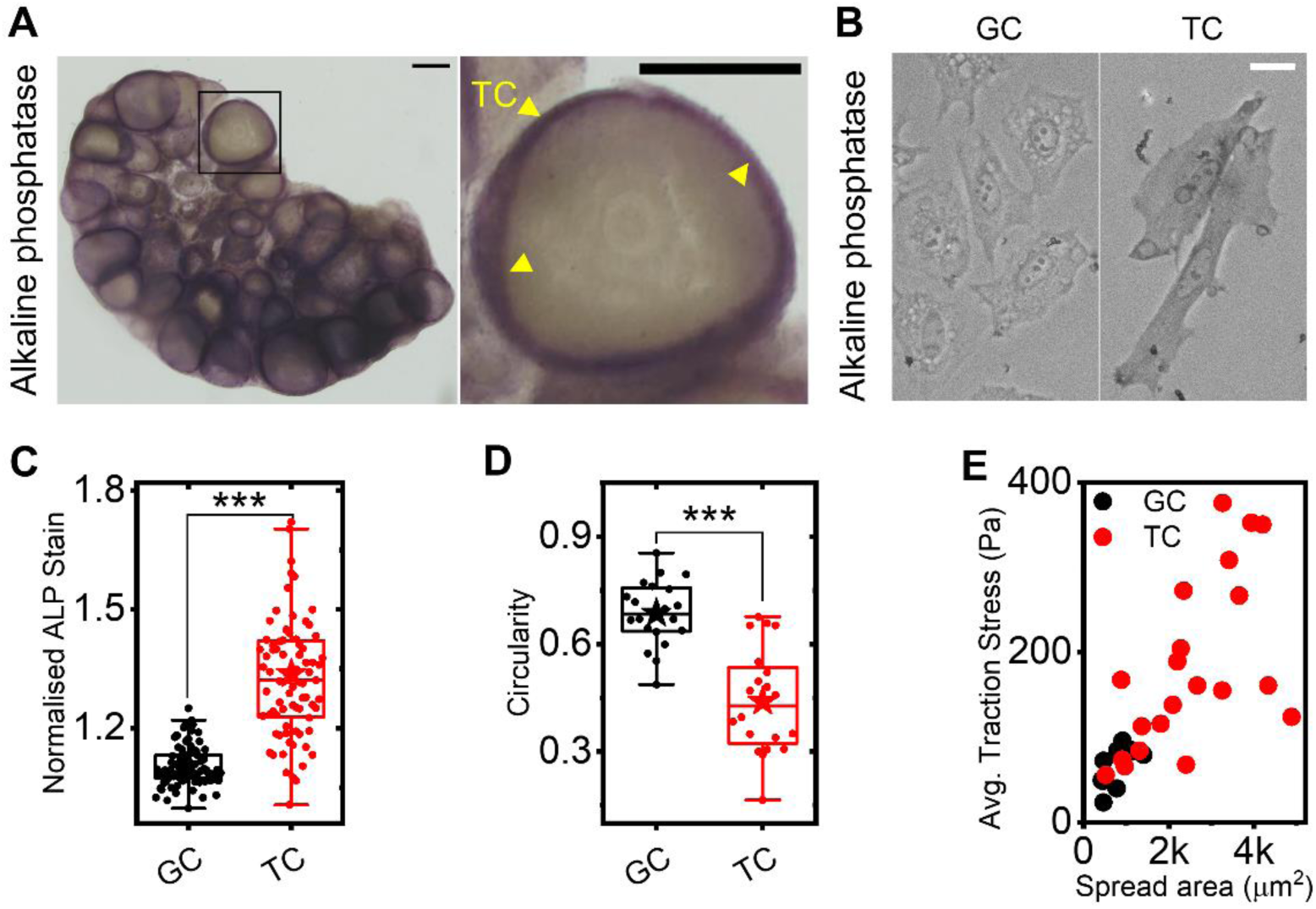
Theca cell purity assessment, related to Figure 1. A) Representative images of alkaline phosphatase staining in ovaries (left) and a zoomed-in view of an outlined follicle (right) showing preferential localisation at the TCs (yellow arrowheads). Scale bar: 200 µm. B) Representative images of alkaline phosphatase staining on primary GCs and TCs cultured *in vitro*. Scale bar: 20 µm. C) Boxplots of alkaline phosphatase intensity for GCs and TCs. D) Boxplots of cell circularity for GCs and TCs. Circularity of 1.0 indicates perfect circular shape. E) Scatter plot of average traction stress of TCs and GCs against their spread area. Significance was determined by Mann-Whitney U test. *** p < 0.001.

**Figure S3:**
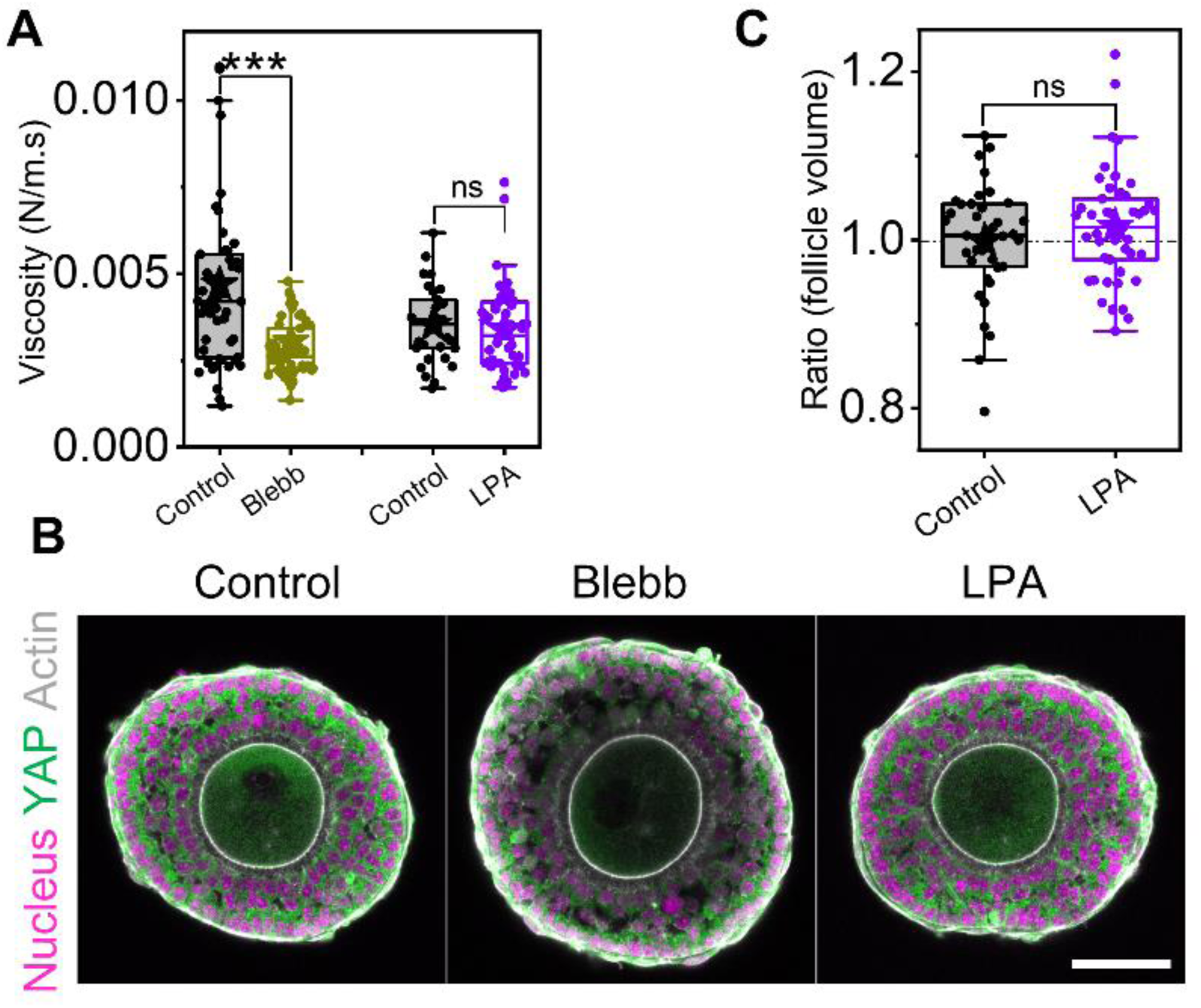
Enhanced TC contractility by LPA has minimal impact on follicle viscosity and size, related to Figure 3. A) Boxplot showing viscosity extracted from AFM indentations in different conditions. N = 5, n = 51 (control), 55 (Blebb); N = 2, n = 31 (control), 51 (LPA) follicles. B) Representative images of isolated follicles labelled with DAPI (nucleus, magenta), Phalloidin (actin, grey), and immuno-stained with YAP (green) in control, Blebb, and LPA-treated samples. Scale bar: 50 µm. C) Boxplots of ratio (follicle volume) in control and LPA-treated samples (30 mins). N = 2, n = 37 (control), 45 (LPA) follicles. Significance was determined by Mann-Whitney U test. ns: p > 0.05; *** p < 0.001.

**Figure S4:**
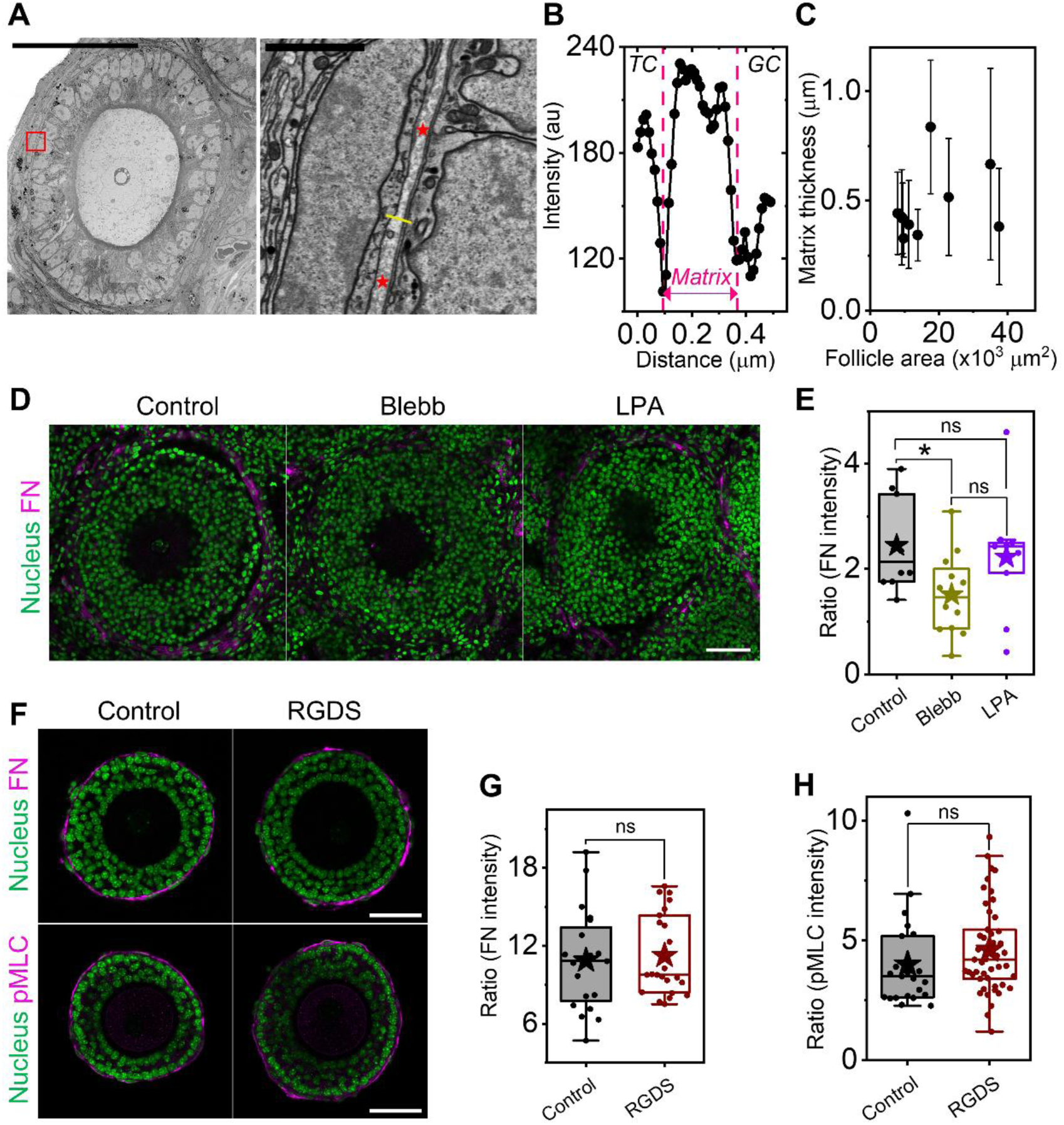
Actomyosin perturbation impacts FN assembly, but disruption of FN-TC coupling does not affect contractility, related to Figure 4. A) Representative SEM images of follicles in an ovarian slice (left, scale bar: 50 µm) and respective zoomed-in sections (right, scale bar: 2 µm) of the red box. Red asterisks mark the matrix between BM and basal TCs. B) Plot profile of the yellow line marked on A. The width of the matrix is marked in magenta. C) Scatter plot of average matrix thickness against follicle area. Error bar represents standard deviation. N = 12 follicles, n = 50 line-scans each. D) Representative images of tissue slices labelled with DAPI (nucleus, green) and immuno-stained with FN (magenta) in control, Blebb, and LPA-treated samples. Scale bar: 50 µm. E) Corresponding boxplots of FN expression of the TC shell in various conditions. N = 1, n = 8-10 follicles. F) Representative images of isolated follicles in control and RGDS-treated samples stained with DAPI (nucleus, green) and immuno-stained with FN (magenta, top) or pMLC (magenta, bottom). Scale bar: 50 µm. G) Corresponding boxplots of ratio (FN intensity) at the TC shell in the two conditions. N = 2, n = 20 (control), 23 (RGDS) follicles. H) Corresponding boxplots of ratio (pMLC intensity) at the TC shell in the two conditions. N = 2, n = 23 (control), 47 (RGDS) follicles. Significance was determined by Mann-Whitney U test. ns: p > 0.05; * p < 0.05.

**Figure S5:**
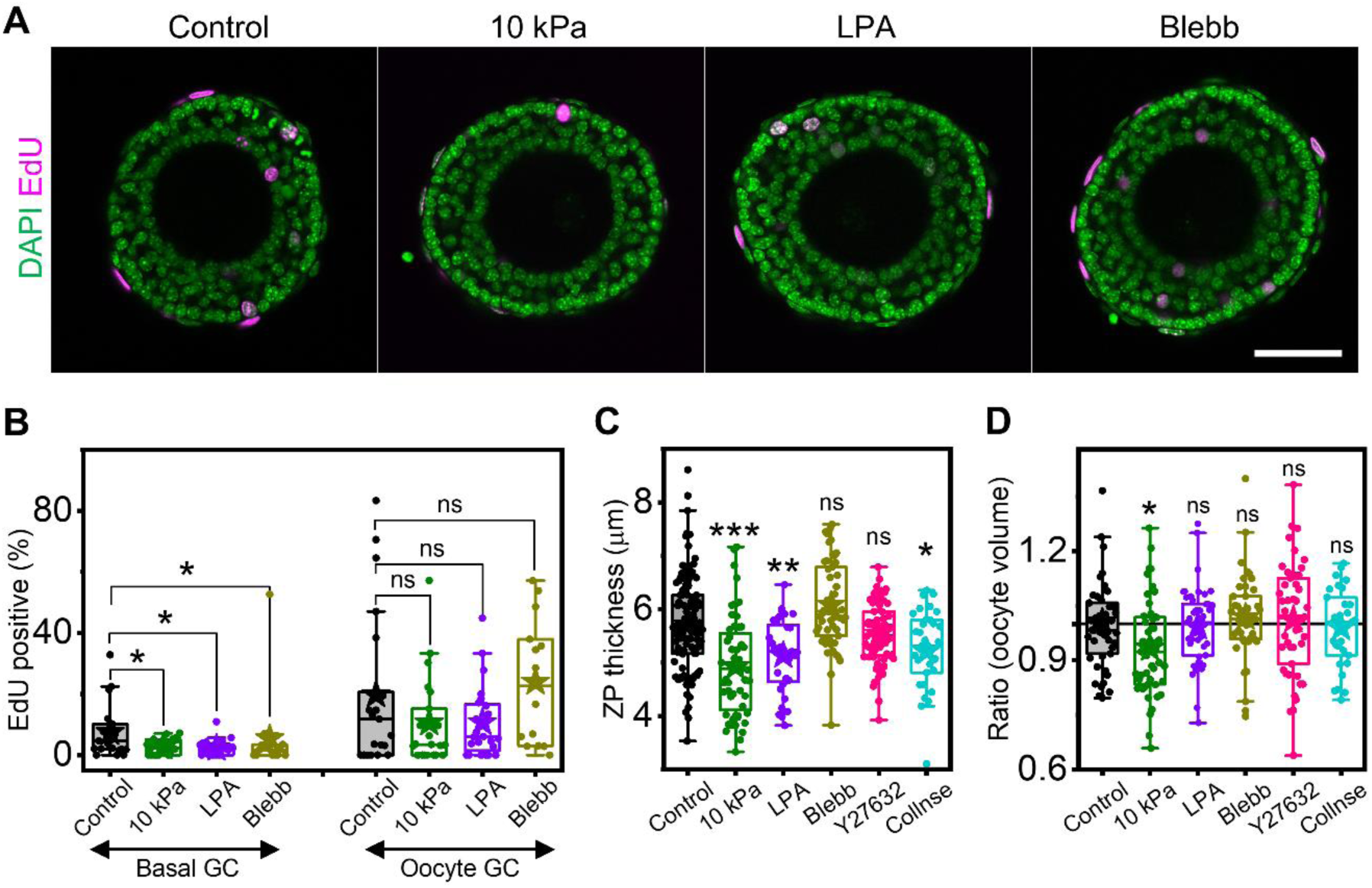
Impact of transient mechanical perturbations on GC proliferation, ZP thickness and oocyte volume, related to Figures 5-6. A) Representative images of DAPI (green) and EdU (magenta) stained isolated follicles in control,10 kPa, LPA, and Blebb-treated samples. Scale bar: 50 µm. B) Corresponding boxplots of EdU-positive basal and oocyte GCs under various mechanical perturbations. N = 2, n = 20 follicles. C) Boxplots of zona pellucida thickness under various mechanical perturbations. N = 4, n = 28-56 follicles in each condition. D) Boxplots of ratio (oocyte volume) under various mechanical perturbations. N = 3, n = 20-35 follicles in each condition. Significance was determined by Mann-Whitney U test. ns: p > 0.05; * p < 0.05; ** p < 0.01; *** p < 0.001.

**Figure S6:**
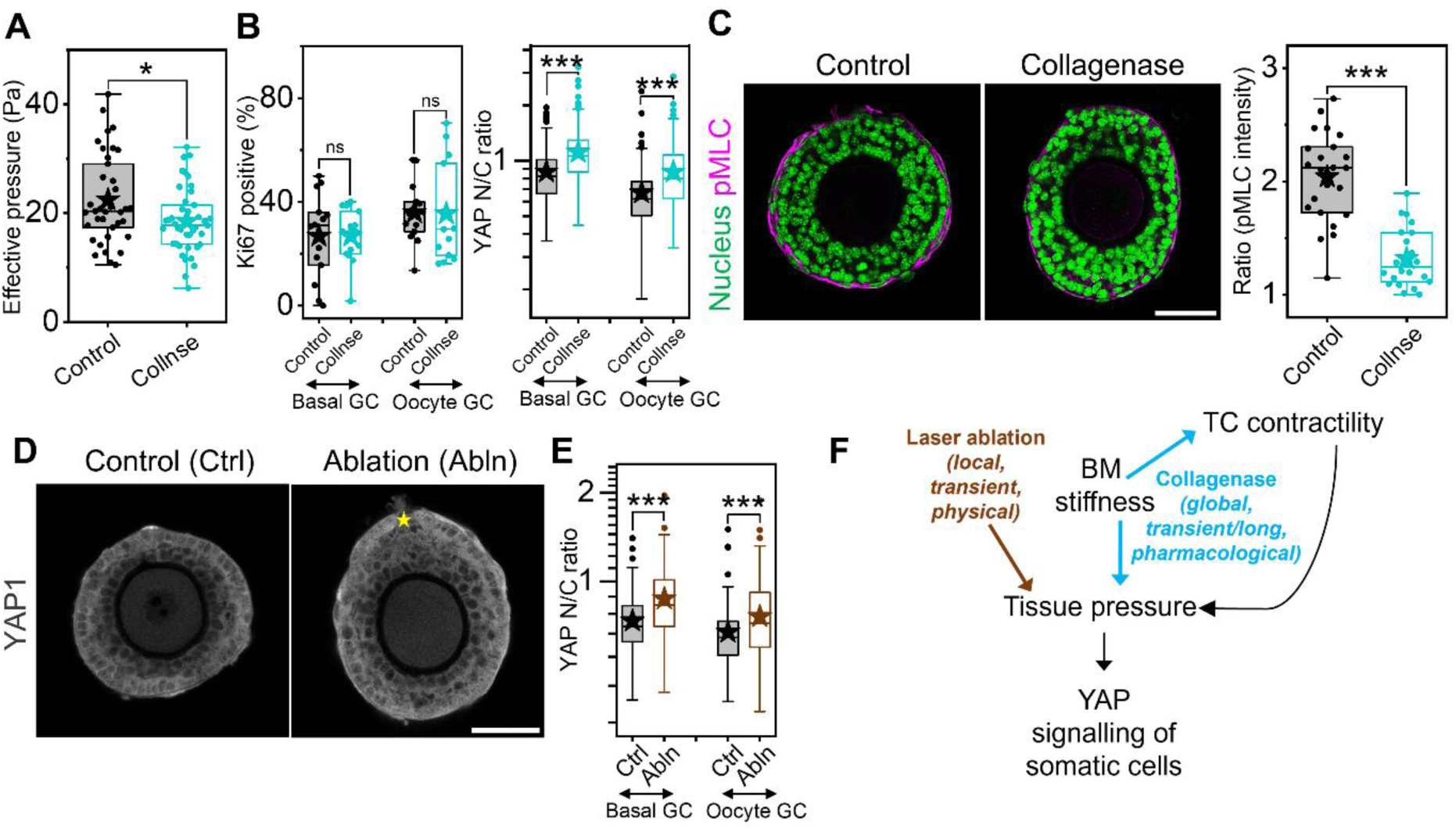
Physical and pharmacological perturbation of tissue pressure affects YAP signalling of granulosa cells, related to Figure 6. A) Boxplots for effective pressure measured by AFM in control and collagenase treated (0.2 mg/ml, 30 mins) follicles. N = 2, n = 37 (control), 43 (collagenase) follicles. B) Boxplots of Ki67^+^ percentage and YAP N/C ratios of basal and oocyte GCs in control and collagenase-treated follicles (0.1 mg/ml, 2 hours). N = 2, n = 18 follicles. C) Left: Representative images showing isolated secondary follicles stained with DAPI (green) and immunolabelled with pMLC (magenta) in different conditions. Scale bar: 50 µm. Right: Boxplots for ratio (pMLC intensity) at TC shell in control and collagenase-treated follicles (0.2 mg/ml, 30 mins). N = 2, n = 20 follicles. D) Representative images of control and laser-ablated follicles immunolabelled with YAP (grey). Yellow asterisk marks the point of ablation. Scale bar: 50 µm. E) Boxplots of YAP N/C ratio of basal and oocyte GCs in different conditions. N = 2, n = 18 (control), 35 (ablation) follicles. F) Schematic representing how manipulation of tissue pressure affects YAP signalling of granulosa cells. Significance was determined by Mann-Whitney U test. ns: p > 0.05; * p < 0.05; *** p < 0.001.

**Figure S7:**
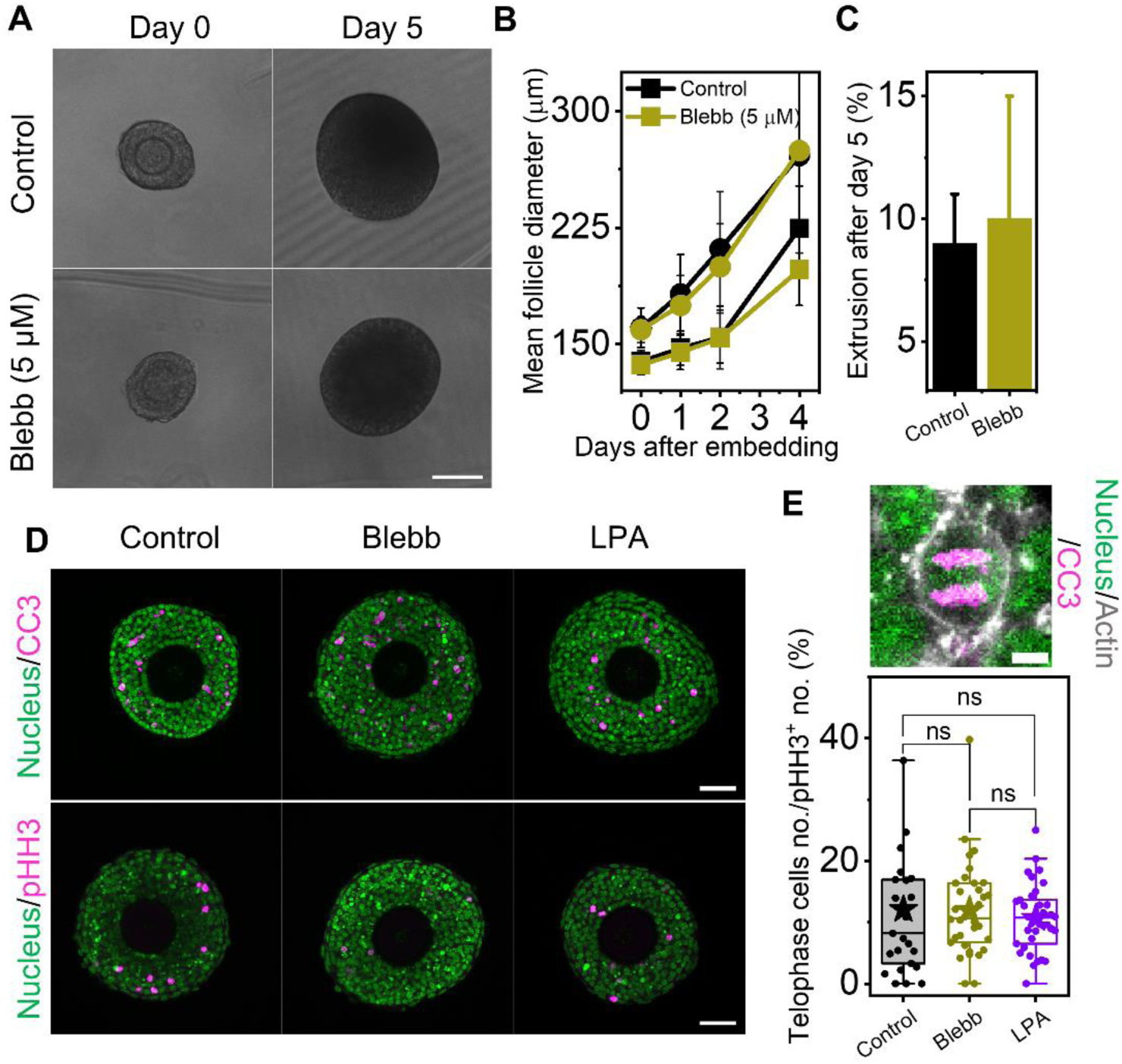
Tissue pressure does not affect apoptosis, proliferation and cytokinesis of GCs in 3D cultures, related to Figure 7. A) Representative images of follicles in control and Blebb (5 µM) conditions at day 0 and day 5 of culture. Scale bar: 100 µm. B) Plot of follicle diameters in the two conditions. N = 3; n = 46 (control), 56 (Blebb) follicles. C) Percentage of extrusion events in the two conditions. Bars represents the average rupture events within an experiment. Error bars represent standard deviation. D) Representative images of 3D-cultured follicles in control, Blebb (20 µM), and LPA-treated conditions, labelled with DAPI (green) and immune-stained with cleaved caspase 3 (CC3, top row) and phospho-histone H3 (pHH3, bottom row) in magenta. Scale bar: 50 µm. E) Top: Representative image showing a cell marked in DAPI (nucleus, green), Phalloidin (actin, grey), and immune-stained with pHH3 (magenta) at telophase in a control follicle. Scale bar: 5 µm. Bottom: Boxplot of telophase cells normalised against the total number of mitotic cells in control, Blebb, and LPA-treated conditions. Significance was determined by Mann-Whitney U test. ns: p > 0.05.

